# A new role of low barrier hydrogen bond in mediating protein stability by small molecules

**DOI:** 10.1101/2021.05.09.443293

**Authors:** Jianhong Yang, Yong Li, Qiang Qiu, Ruihan Wang, Wei Yan, Yamei Yu, Lu Niu, Heying Pei, Haoche Wei, Liang Ouyang, Haoyu Ye, Dingguo Xu, Yuquan Wei, Qiang Chen, Lijuan Chen

**Author notes:** Correspondence should be addressed to L.C. Additional corresponding authors: Q.C. These authors equally contribute to this work (Jianhong Yang, Yong Li, Qiang Qiu, Ruihan Wang).

## Abstract

Low barrier hydrogen bond (LBHB) is a special type of hydrogen bond which occurs where two heteroatoms with similar p*Ka* values share a single proton resulting in an unusually strong and short hydrogen bond. LBHBs in protein play important roles in enzyme catalysis and maintaining protein structural integrity but its other biochemical roles are unknown. Here we report a novel function of LBHB in selectively inducing tubulin protein degradation. A tubulin inhibitor, 3-(3-Phenoxybenzyl) amino-β-carboline (PAC), promotes selective degradation of αβ-tubulin heterodimers by binding to the colchicine site of β-tubulin. Biochemical studies have revealed that PAC specifically destabilizes tubulin, making it prone to aggregation that then predisposes it to ubiquitinylation and then degradation. Structural activity analyses have indicated that the destabilization is mediated by a single hydrogen bond formed between the pyridine nitrogen of PAC and βGlu198, which is identified as a LBHB. In contrast, another two tubulin inhibitors only forming normal hydrogen bonds with βGlu198 exhibit no degradation effect. Thus, the LBHB accounts for the degradation. Most importantly, we screened for compounds capable of forming LBHB with βGlu198 and demonstrated that BML284, a Wnt signaling activator, also promotes tubulin heterodimers degradation in a PAC-like manner as expected. Our study has identified a novel approach for designing tubulin degraders, providing a unique example of LBHB function and suggests that designing small molecules to form LBHBs with protein residues resulting in the highly specific degradation of a target protein could be a new strategy for drug development.

## 1. Introduction

Hydrogen bond was firstly discovered almost a century ago and it played an important role in physics, chemistry and biology [1]. In life science, it is extremely important for various biological processes, including the formation of DNA double helix structure, protein secondary structure, protein folding and stability, enzyme catalysis, drugs binding and molecular recognition [2-7]. A special hydrogen bond named low barrier hydrogen bond (LBHB) was first proposed in 1990s [8, 9]. LBHB is distinguished from conventional hydrogen bond by equally proton sharing between two heteroatoms [10]. These two heteroatoms usually have similar p*Ka* values and form an unusually strong and short (≤2.65 Å for O-N pairs) hydrogen bond [3, 11, 12]. The proton of an LBHB lies in the middle of the two heteroatoms and precise location of the proton by experiment could provide direct evidence of existence of LBHB [13]. To locate the proton of an LBHB in a protein can only be done by ultra-high-resolution (1.0 Å or less) X-ray or neutron crystallography while it is technically difficult for most proteins [13]. Moreover, theoretical methods such as quantum mechanics/molecular mechanics (QM/MM) calculation of potential-energy curves of proton has been widely used to predict LBHB [13-15]. Formation of an LBHB can supply 10-20 kcal/mol energy which can facilitate difficult biochemical reaction [9]. Thus, LBHB is regarded to be responsible for catalytic efficiencies of several enzymes including aspartate aminotransferase, ketosteroid isomerase, citrate synthase, serine proteases and aminoglycoside-N3-acetyltransferase Via [16-20]. Besides, LBHBs are needed to maintain protein structural integrity in proteins including rhamnogalacturonan acetylesterase D192N mutant and human transketolase [21, 22]. However, other biochemical roles of LBHBs are rarely reported.

LBHBs is also formed between small-molecule ligand and proteins, such as human transketolase and periplasmic phosphate-binding proteins, to mediate ligand binding and selectivity, implying LBHBs have the potential to be used for high selective drug design [22, 23]. The intermolecular hydrogen bond between carboxylic acid and pyridine was extensively studied and both experimental and theoretical calculation evidences revealed this intermolecular hydrogen could be an LBHB [24-26]. Most recently, an LBHB was found between pyrimidyl of thiamine and carboxyl of glu366 in human transketolase through ultra-high-resolution X-ray crystallography [22]. Thus, it is possible that an LBHB could exist between inhibitors (containing pyridine or pyridine-like function group) and its target protein and may have special biochemical functions.

A synthetic compound, 3-(3-Phenoxybenzyl)amino-β-carboline (PAC, Fig 1a), belongs to the class of known anti-cancer natural products [27] denoted β-carboline derivatives [28-30]. PAC is a potent tubulin inhibitor with nanomolar IC_50_ values in several tumor cells [30]. Tubulin is a validated target for a wide range of tumors and several drugs, including paclitaxel, eribulin and vinca alkoloids, that inhibit the polymerization or depolymerization of tubulin are in clinical use [31, 32]. We have determined that PAC has a novel mode of action beyond simple inhibition of tubulin polymerization. We show that PAC when given in relevant doses results in the specific degradation of tubulin by the ubiquitin pathway. Our data indicate that ubiquitinylation is a response to destabilization and aggregation of tubulin. Structural activity analysis reveals that an LBHB between pyrimidyl of PAC and carboxyl of βGlu198 underpins the degradation. While another two compounds forming normal hydrogen bonds with βGlu198 exhibited no tubulin degradation effect. We screened for other compounds that can form LBHB with βGlu198 and identified that BML284, a Wnt signaling activator also promoted tubulin heterodimers degradation in a PAC like manner.

**Figure 1:**
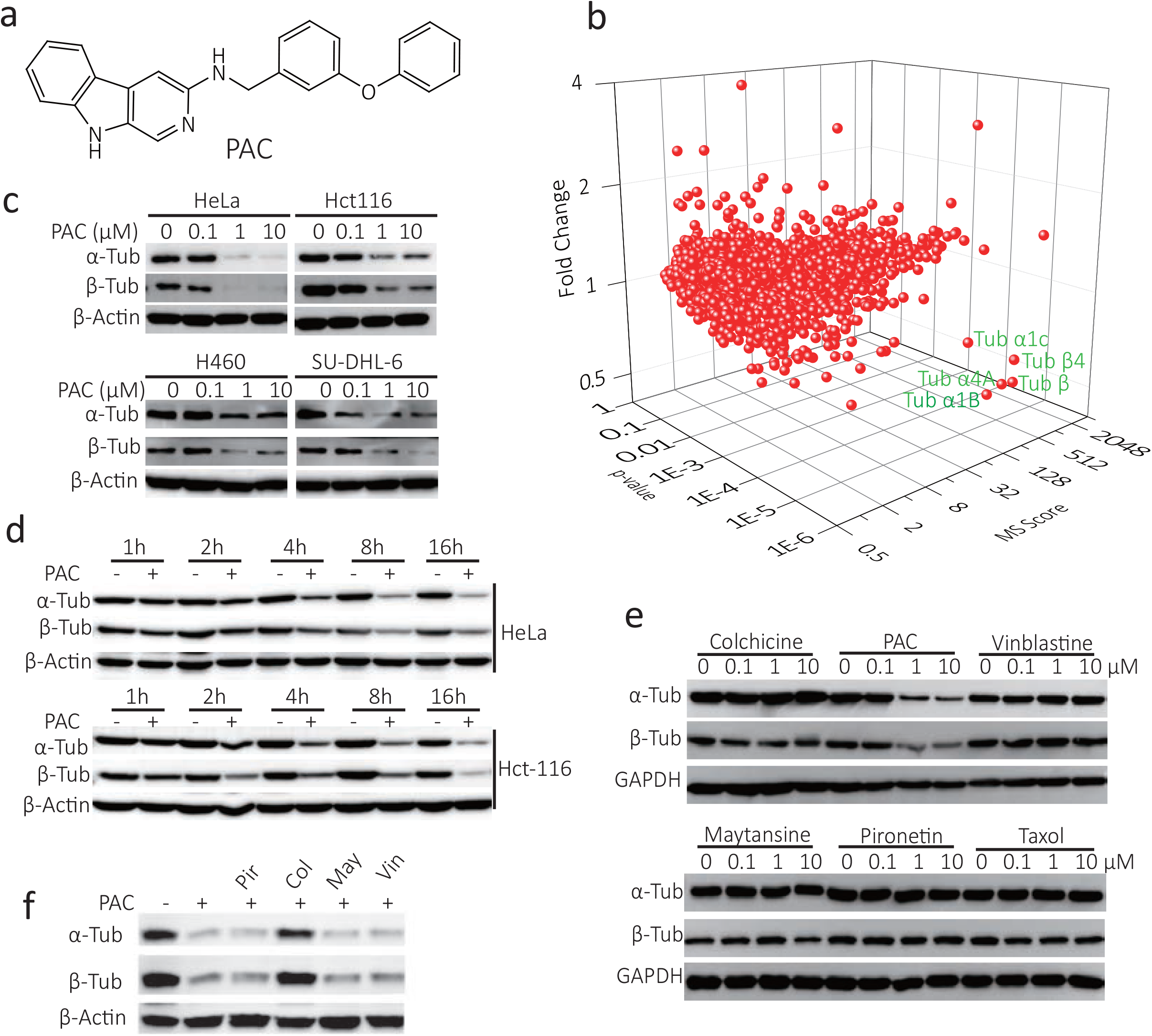
PAC selectively downregulates the protein level of αβ-tubulin heterodimers. (**a**) Chemical structure of PAC. (**b**) Proteome-wide analysis of HeLa cells treated with 1 μM PAC or DMSO alone for 6 hours using tandem mass tag quantitative proteomics. Each point represents the ratio of mean protein levels of three biological repetition in PAC-versus DMSO-treated cells. (**c**) Immunoblot analysis of α- and β-tubulin protein levels in HeLa, Hct116, H460 and SU-DHL-6 cells treated with the indicated concentrations of PAC for 24 h. β-Actin was used as loading control. Results are representative of three independent experiments. (**d**) Time course of α- and β-tubulin protein levels in HeLa and Hct116 cells treated with 1 μM PAC or DMSO. β-Actin was used as the loading control. Immunoblot results are representative of three independent experiments. (**e**) Comparing the tubulin degradation effect of PAC with conventional tubulin inhibitors. HeLa cells were treated with different concentrations of PAC, colchicine, vinblastine, maytansine, pironetin and taxol for 16hours, then α- and β-tubulin protein levels were detected by immunoblot. Results are representative of two independent experiments. (**f**) Inhibition of αβ-tubulin heterodimers degradation by tubulin inhibitors. HeLa cells were treated with or without 1μM pironetin, colchicine, maytansine and vinblastine for 1hour before treated with or without 1μM PAC for 16hours. Then cell α- and β-tubulin protein levels were monitored by immunoblot analysis. Results are representative of three independent experiments.

Our study demonstrated a new role of LBHB in mediating protein stability by small molecules, identified a novel approach to design tubulin degraders and suggest a new strategy for drug development, designing small molecules to form LBHBs with protein residues resulting in the highly specific degradation of a target protein. Also, the interaction between the carboxylic acid and pyridine or pyridine-like function group could be used as a model to design drugs forming LBHB with target protein.

## 2. Results

### 2.1 PAC selectively downregulates the protein level of tubulin heterodimers

PAC (Fig.1a) showed high anticancer activity *in vitro* [33], and is identified as a tubulin inhibitor [30]. PAC induced obvious G2/M phase cell cycle arrest on human cervical adenocarcinoma cell line HeLa and human colon colorectal carcinoma cell line Hct116 (**Fig.S1a**) and caspase-dependent apoptosis on HeLa cells (**Fig.S1b)**. PAC significantly and dose-dependently inhibited tumor growth of both human large cell lung carcinoma cell line H460 and Hct116 xenografts (**Fig.S1c** and **S1d**). To illuminate the cellular effect of PAC in an early time point, we employed a tandem mass tag (TMT) quantitative proteomic analysis of six-hour PAC treated HeLa cells, and demonstrated that PAC could specifically downregulate the protein level of α- and β-tubulin and their subtypes (**Fig.1b**). The biological processes and cellular components analysis of the quantitative proteomic data showed that the mitotic cell cycle was the most effected biological process and cytoskeleton was the most effected cellular component in response to PAC treatment (**Fig.S2a** and **S2b**). These results were consistent with tubulin reduction since tubulin is the main component of cytoskeleton and plays an important role in mitosis [34]. Immunoblot analysis similarly exhibited clear reduction of α- and β-protein in HeLa, Hct116, H460 and human B cell lymphoma cell SU-DHL-6 in a dose (**Fig.1c**) and time dependent manner for HeLa and Hct116 cells (**Fig 1d**). A qPCR assay confirmed that the drop in level of αβ*-*tubulin proteins is post transcriptional, since mRNA levels of αβ*-*tubulin in HeLa and Hct116 showed no such time dependence (**Fig. S3a** and **S3b**). We also found PAC administration reduced the protein level of tubulin in tumor tissue of an Hct116 xenograft model (**Fig.S3c**). *In vitro* tubulin polymerization assay showed that PAC directly binds to purified tubulin and inhibits tubulin polymerization in a concentration dependent manner (**Fig.S3d**). Of all the widely known tubulin-binding agents we have tested, only PAC could induce the degradation of tubulin (**Fig.1e**). To identify the binding site of PAC on tubulin and whether the binding of PAC to tubulin accounts for the degradation effect, we carried out competition assays and found that colchicine, but not vinblastine, maytansine or pironetin inhibit PAC-induced tubulin degradation, suggesting that PAC exerts it effect by binding to the colchicine site (**Fig.1f**). All these indicated PAC induces selective tubulin degradation by binding to the colchicine site.

### 2.2 PAC specifically destabilizes tubulin making it prone to aggregation that then predisposes it to ubiquitinylation

HeLa cells were pretreated with various inhibitors, including lysosome inhibitor NH_4_Cl, proteasome inhibitor MG132, autophagy inhibitor 3-MA. Only the proteasome inhibitor MG132 completely blocked PAC-induced tubulin degradation (**Fig.2a** and **S4a**), suggesting proteasome pathway is responsible for the tubulin degradation. Immunoblot analysis of immunoprecipitated α- and β-tubulin by ubiquitin antibody showed that PAC treatment has significantly increased the ubiquitinylation level of α- and β-tubulin (**Fig.S4b**), whereas no such obvious effect on the general ubiquitinylation level of the total proteins (**Fig.S4c**) was observed. PYR-41, a specific inhibitor for ubiquitin-activating enzyme E1, completely inhibited PAC-induced tubulin degradation (**Fig.2a**), suggesting PAC induced tubulin degradation relied on ubiquitinylation. Cells pretreated with MG132 (or PYR-41) possessed aggregates of tubulin in cytoplasm after PAC treatment (**Fig.2b**). Size-exclusion chromatography also revealed that PAC treatment resulted in purified tubulin aggregation *in vitro* (**Fig.2c**), which was further confirmed by transmission electron microscope (TEM) with lots of different size spherical tubulin aggregates observed (**Fig.2d**). We speculated that PAC might promote denaturation of tubulin and ubiquitinylation arise from normal protein clearance. Using a Thiol Fluorescent Detection Kit the denaturation of both tubulin and bovine serum albumin (BSA) were studied. While PAC showed no denaturation effect on BSA, it promoted denaturation of tubulin at low concentrations, an effect that was inhibited by pretreatment with colchicine (**Fig.2e**), confirming PAC induces selective denaturation of tubulin *in vitro*. Differential scanning fluorimetry results indicated tubulin treated with PAC showed obvious lower transition midpoint (Tm) whereas colchicine had no effect (**Fig.2f**), demonstrating PAC decreases the stability of tubulin *in vitro*. We conclude then that PAC specifically destabilizes tubulin making it prone to aggregation that then predisposes it to ubiquitinylation.

**Figure 2:**
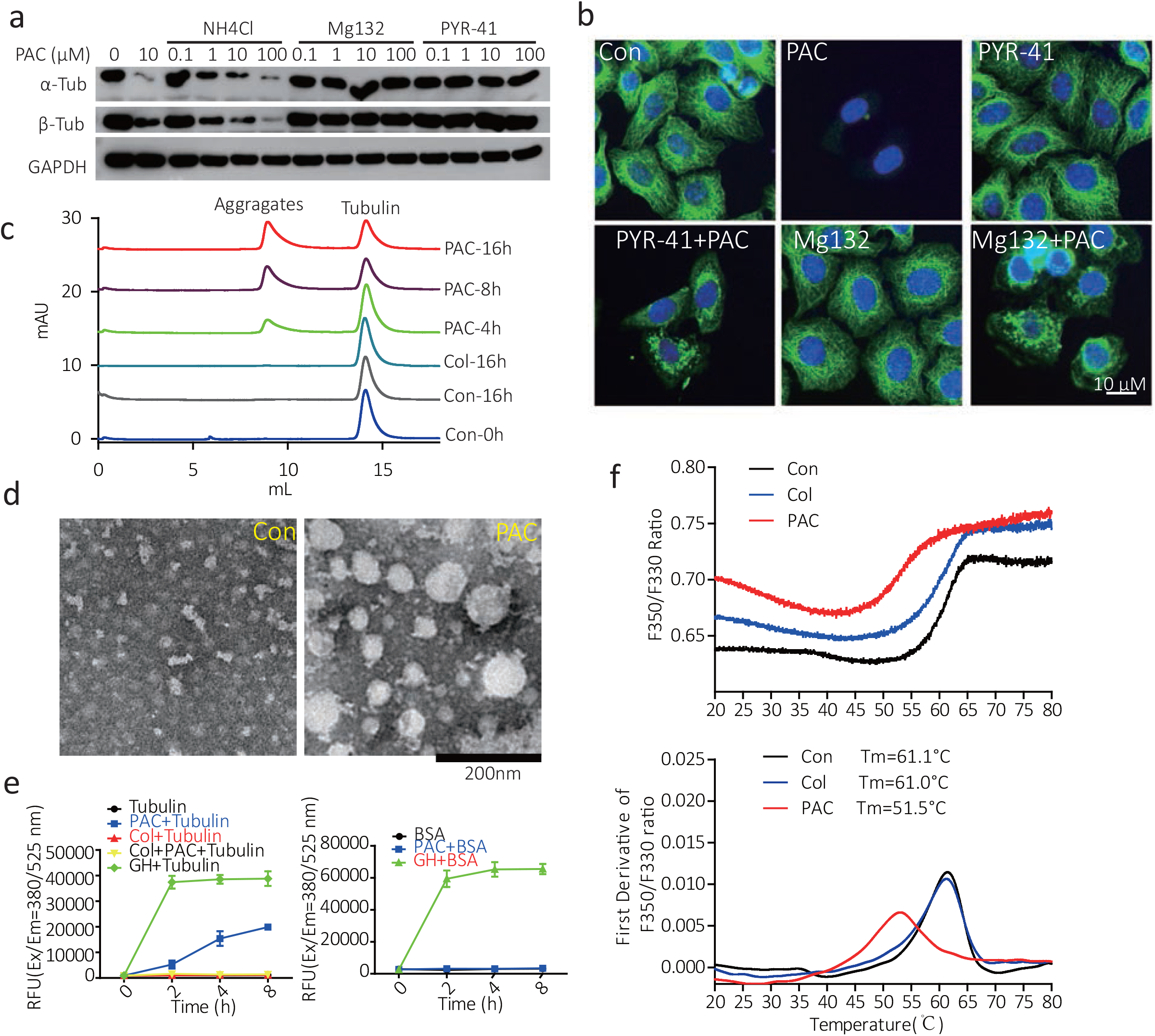
PAC promotes tubulin destabilization and aggregation. (**a**) Proteasome and ubiquitin inhibitors rescue PAC-induced tubulin degradation. HeLa cells were pretreated with 20 mM NH4Cl, 20 μM Mg132 or 20 μM PYR-41 for one hour before the addition of PAC. Results are representative of three independent experiments. (**b**) PAC induced tubulin aggregation in cytoplasm. HeLa cells were pretreated with Mg132 or PYR-41 for 1 hours before treated with PAC for 4 hours. The tubulin morphology was monitored by immunofluorescence. Results are representative of three independent experiments. (**c**) Gel filtration assays detection of tubulin aggregation. Purified tubulin was dissolved to 3μM in PEM buffer at room temperature before incubated with different compounds for different times. Aer incubation, samples were analyzed by a gel filtration assay. Results are representative of two independent experiments. (**d**) Transmission electron microscopy (TEM) detection of tubulin. Purified tubulin (10 μM) was incubated with or without 20μM PAC for 4 hours at room temperature, the images were taken by TEM. Results are representative of two independent experiments. (**e**) PAC promotes purified tubulin denaturation. Tubulin (1μM) was incubated with guanidine hydrochloride(GH)or other compounds or DMSO for 2, 4 or 8hours at room temperature. Then master reaction mix was added to the samples and incubated for another 30min. At last the fluorescent adduct were detected by a Biotech Gen5 spectrophotometer (λex = 380/λem = 525 nm). Data are shown as the mean ± SD (n = 3). (**f**) Thermal unfolding curves of DMSO, PAC (30 μM) or colchicine (30 μM) treated purified tubulin(3 μM)by a differential scanning fluorimetry (DSF) method. Plots of the fuorescence F350/F330 ratio (up) and the first derivative of F350/F330 ratio (low) are shown. Results are representative of three independent experiments.

### 2.3 Crystal structure of tubulin-PAC complex

Crystals of a protein complex composed of two αβ-tubulin heterodimers, the stathmin-like protein RB3 and tubulin tyrosine ligase (T2R-TTL) [35] were soaked with PAC and the resulting complex determined to 2.65 Å resolution (**Fig.3a**). Details of the data collection and refinement statistics are summarized in **Supplementary Table 1**. The Fo–Fc difference electron density identified there was a PAC molecule on each of β1-tubulin and β2 tubulin (**Fig.3b**). Our previous structure-based pharmacophore for colchicine binding site inhibitors (CBSIs) suggest there are five recognition centers: three hydrophobic centers (I, II and III) and two hydrogen bond centers (IV and V) [36]. In this tubulin-PAC complex, PAC occupied hydrophobic centers I and II where it makes extensive van der Waal contacts (**Fig.3c**). PAC forms hydrogen bonds with the side chains of βE198 and βY200 in hydrogen bond center IV (**Fig.3c**). Compared to the colchicine complex (PDB:4O2B) [37], PAC was located deeper in the β subunit and made no interaction with α subunit consequently the atoms of PAC and colchicine show only little overlap (**Fig.S5a**). Comparison between tubulin-PAC and tubulin-colchicine complex structures showed that the binding did not affect the global conformation of tubulin nor of the T2R complex. The root mean square deviation (RMSD) for 1,967 Cα atoms between tubulin-PAC and tubulin-colchicine complexes is 0.4 Å.

**Figure 3:**
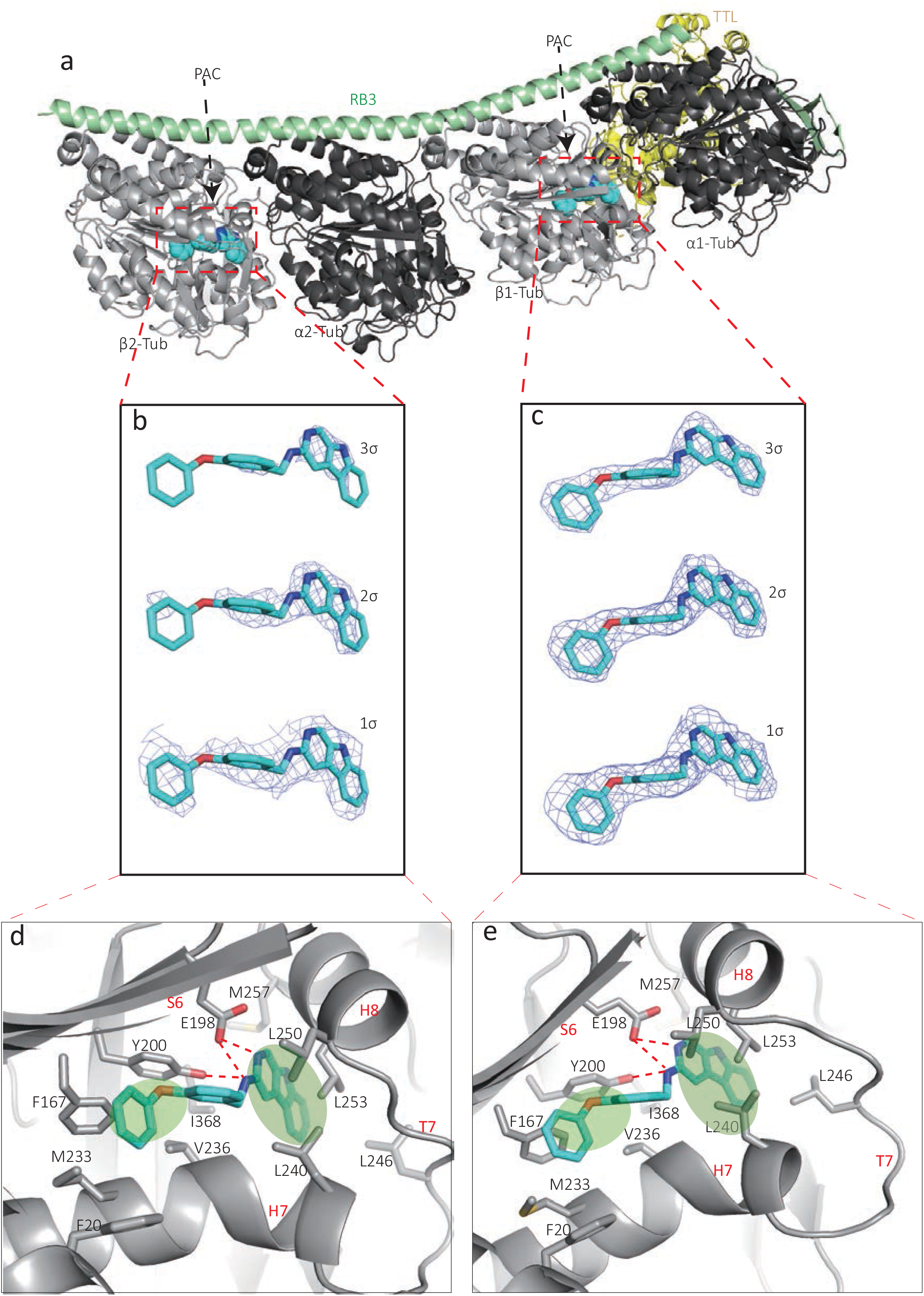
Crystal structure of PAC-tubulin complex. (**a**) Overall structure of tubulin-PAC complex. The RB3-SLD is colored green, TTL is yellow, α-tubulin is black, β-tubulin is grey, and PAC is cyan. PAC is shown as spheres. (**b, c**) Electron densities of PAC on (**b**) β2-tubulin or (**c**) β1-tubulin. The Fo-Fc omit map is colored light blue and contoured at 3δ,2δ and 1δ. (**d, e**) Interactions between PAC and (**d**) β2-tubulin and (**e**) β1-tubulin. Residues that make interactions with PAC are shown as scks and are labeled. The hydrophobic centers are indicated by green semitransparent ovals (small: center I; big: center II). Hydrogen bonds are drawn with red dashed lines.

### 2.4 The pyridine nitrogen of PAC and Glu198 of β-tubulin mediate the tubulin degradation

Informed by the crystal structure PAC-tubulin complex, we synthetized a series of PAC derivatives to identify which functional group of PAC is responsible for its tubulin degradation ability (**Fig.4a**). Compounds **9, 10, 11, 14** and **15** showed anti-proliferation and promoted tubulin degradation at higher concentrations (**Fig.4b, 4c** and **S5b**). For ease of discussion, we decomposed PAC into three rings (**Fig.4a**). Since compound **9** lacked the **C** ring of PAC, we conclude it is not essential. The activity of compounds **10** and **11** exhibited that the **B** ring can be substituted by aliphatic ring or chain. The binding mode of **9**– and **11**–tubulin complexes were similar to PAC (**Fig. S5c**). Compounds **14** and **15** revealed that even the carboline ring (A ring) is not critical (**Fig 4d, 4e** and **S3b**). Compounds **20** and **24** were synthesized to probe the importance of N-2 and -NH-group at C-3 position. Compound **20** lost both anti-proliferation and tubulin degradation activity while compound **24** retained both (**Fig. 4b, 4c** and **S5b**). We concluded that N-2 on the carboline ring is the key functional group contributing to tubulin degradation. The crystal structure of PAC-tubulin complex demonstrated that only βGlu198 and βTyr200 make polar interactions with N-2 of PAC. PAC addition promoted wild-type like degradation of Y200F β-tubulins but had no effect on E198G, E198Q or E198D β-tubulins (**Fig.4d**). EBI assay revealed that PAC no longer bound to the colchicine site of β-tubulin mutants βE198G, βE198Q, or βE198D (**Fig.S5d**). We concluded that the hydrogen bond interaction between βGlu198 and the nitrogen atom at 2-position of A ring of PAC triggers tubulin degradation. However, two tubulin inhibitors, nocodazole (Noc) and plinabulin (Pli), adopt similar conformations in β-tubulin to PAC. and also form hydrogen bonds with βGlu198 (**Fig.4e**) [38], they could not cause tubulin degradation effect and block PAC-induced degradation (**Fig.4f**), suggesting that the hydrogen bond between OGlu198 and pyridine nitrogen of PAC may be special.

**Figure 4:**
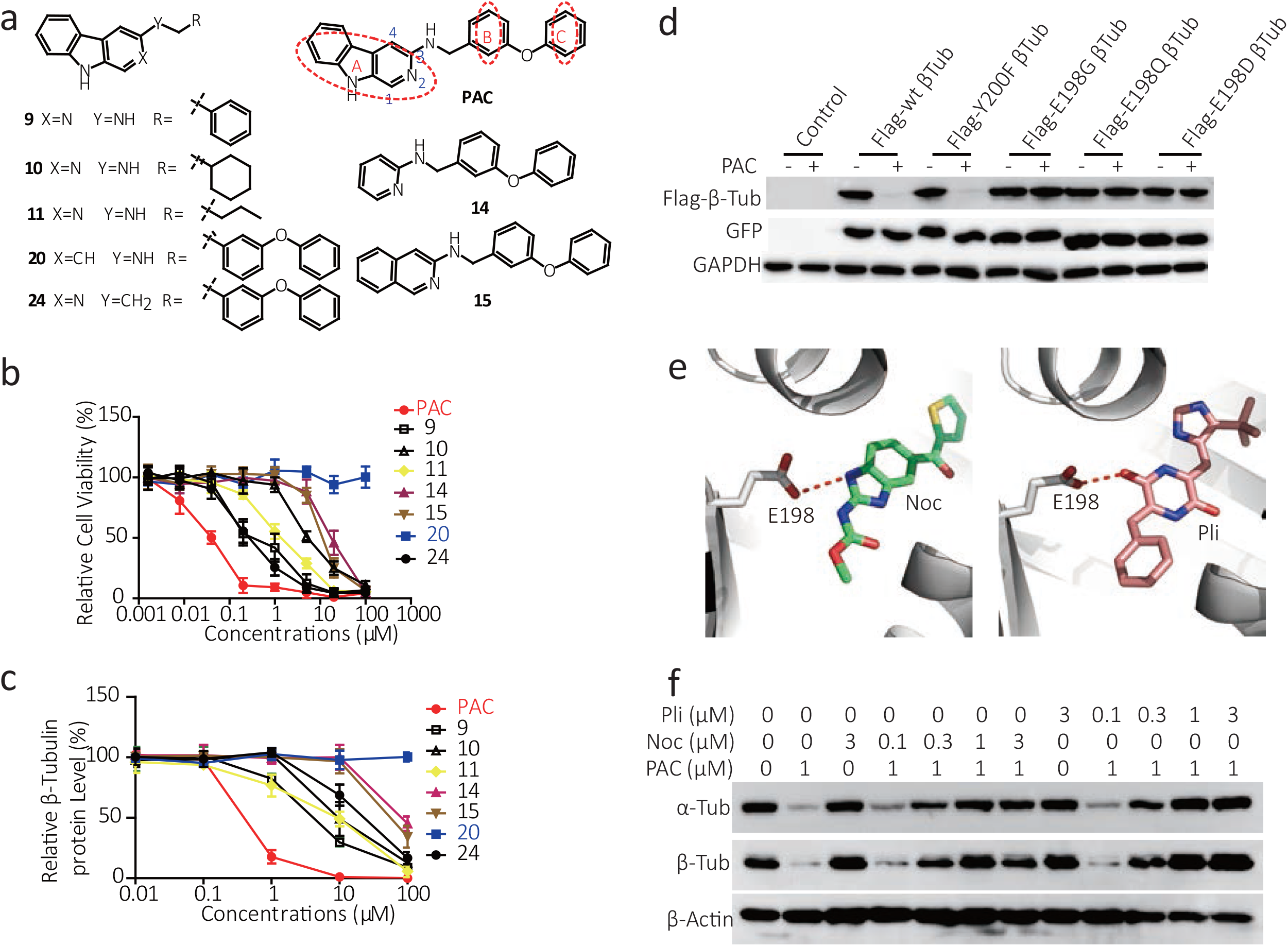
The pyridine nitrogen of PAC and Glu198 of β-tubulin mediate the tubulin degradation. (**a**) Chemical structure of synthec PAC derivatives. (**b**) Growth inhibitory curves of PAC derivatives in HeLa cells. Cells were treated with different concentrations of indicated compound for 48 hours, cell viability were monitored by MTT assay. Data are shown as the mean ± SD (n = 3). (**c**) Tubulin degradation activity of PAC derivatives. HeLa cells were treated with different concentrations of indicated compound for 16hours, the tubulin protein levels were monitored by immunoblot. Densitometric quantification of immunoblot was performed by Image Quant TL software, Data are shown as the mean ± SD (n = 3). (**d**) Effect of PAC on tubulin mutants. MSCV-IRES-GFP vectors expressing both GFP and wild type or mutant Flag-β-tubulin were transfected to HeLa cells. After 24 hours, cells were treated with or without PAC for 16hours. Then the protein level of Flag-tubulin and GFP were detected by immunoblot. GAPDH were used as loading control. Results are representative of three independent experiments. (**e**) Close-up views of the interaction between βGlu198 with nocodazole (Noc) and plinabulin (**f**) Inhibition of tubulin heterodimers degradation by colchicine site tubulin inhibitors. Hela cells were treated with different concentrations nocodazole and plinabulin for one hour before addition of PAC. Results are representative of two independent experiments.

### 2.5 An LBHB between PAC and βGlu198 triggers tubulin degradation

Several previous papers have identified the formation of LBHBs between carboxyl and nitrogen of pyridyl in compounds complexes through chemical calculations, ultralow temperature NMR and cocrystal structure studies [22, 24, 26, 38], which was almost the same pattern with PAC-βGlu198 model. We calculated that the p*K*a value (using Propka 3.1) of βGlu198 to be 9.6, indicating that it is neutral (protonated) under normal conditions. Upon PAC binding, the p*K*a values of βGlu198 and pyridine nitrogen of PAC in the PAC-crystal structure are 2.3 and 2.5 respectively, similar enough to indicate the presence of an LBHB [39]. The interatomic distance between O_Glu198_ and compounds in PAC-tubulin, Compound **9**-tubulin and Compound **11-**tubulin complex is ranged from 2.66 to 2.83 Å (**Fig.5a**). Hydrogen bonds length in X-Ray crystal structure is aligned longer than that in neutron crystal structure. For example, the length of the Glu198-His198 LBHB in X-ray structure of sisomicin bound AAC-Via (6BC7) is 2.84 Å, while in the neutron/X-ray structure (6BBZ) is 2.57 Å [20]. Thus, the length of O_Glu198_-N_PAC_ hydrogen bond could also meet the requirement of an LBHB. Besides, in the QM/MM Optimized structure (more accurate than experimental structure for reaction mechanism study [40-42]) the interatomic distance between O_Glu198_ and N_PAC_ is 2.49 Å (**Fig.S6a**), which is far smaller than the sum of their van-der-Waals radii (2.65 Å for O-N pairs) and meets the need of an LBHB. Using QM/MM calculation, a mature and widely used method for predicting LBHB [13, 22, 43-45], we found the potential energy curve of O_Glu198_-N_PAC_ hydrogen has a double-well character, and the computed proton transfer barrier is 1.77 kcal/mol, which is well under the threshold that typically indicates an LBHB and is under the sum of zero point vibrational energy (ZPVE, 1.46 kcal/mol) and kT (0.596 kcal/mol) (**Fig.5b**) [14]. Besides, QM/MM molecular dynamics simulations clearly showed the proton is able to move freely between βGlu198 and PAC (**Fig.5c-d, S6b-d**). In normal hydrogen bond, the splitting of vibrational energy decreases as the vibration step rises (The |1⟩ to |2⟩ energy ε _21_ is smaller than the |0⟩ to |1⟩ energy ε _10_), while the PAC vibration energy splitting increases with the rise of the vibration step (ε _21 >_ ε _10_)(**Table S2**), resulting a superharmonicity effect, which also suggested an LBHB [46]. In contrast, the potential energy curves and vibration energy splitting decrease (ε _21 <_ ε _10_) of O_Glu198_-N_Noc_ and O_Glu198_-O_Pli_ indicate these two tubulin inhibitors could not form LBHBs (**Fig.5b and Table S2**). All these demonstrated that there is an LBHB formed between O_Glu198_ and N_PAC_ and the LBHB is required for the tubulin degradation effect.

**Figure 5:**
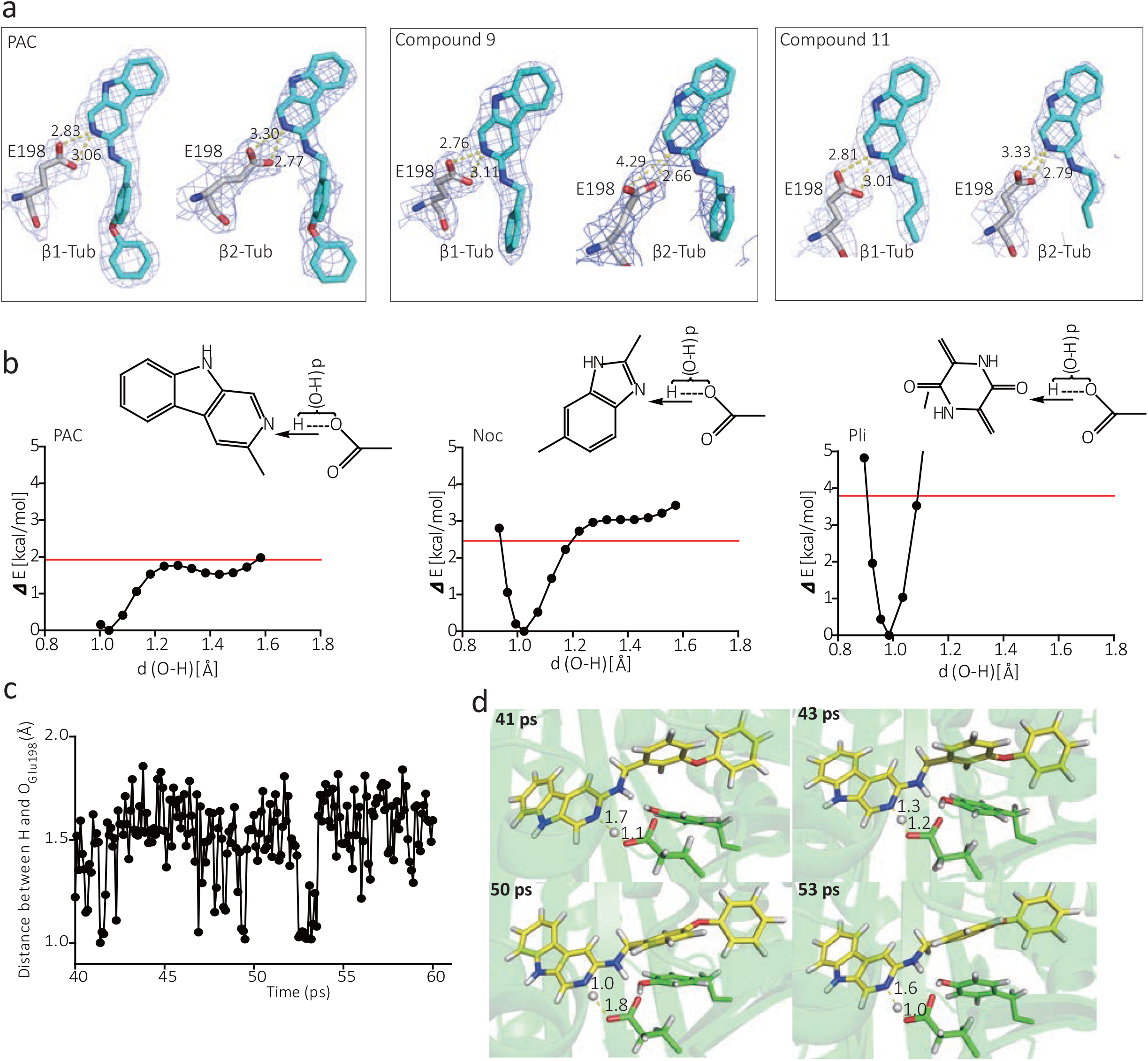
An LBHB between PAC and βGlu198 mediates tubulin degradation. (**a**) Distance between nitrogen of compounds (PAC, compound 9 and compound 11) to βGlu198. The 2Fo-Fc omit map is colored light blue and contoured at 1δ. Compounds and βGlu198 is shown in sticks and colored in cyan and grey respectively. (**b**) Energy profiles along the proton transfer coordinate for H-bond donor-acceptor pairs of the OGlu198-NPAC, OGlu198-NNoc and OGlu198-OPli hydrogen bonds. The red lines are the sum of zero-point vibrational energy and kT (0.596 kcal/mol). (**c**) The distance between proton and O of GLU198 in the 40 to 60 ps QM/MM MD simulaon. (**d**) Snapshots of tubulin-PAC complex along the dynamic simulaon time. For clarity, the water molecules have been removed. The distances between protonated and the O atom/N atom are marked in the figure.

### 2.6 Identification of a new tubulin degrader with the LBHB mediated degradation mechanism

To further confirm of the degradation mechanism and identify effective new tubulin degradation compounds, we screened compounds from lots of tubulin inhibitors with molecular docking and QM/MM calculation and found BML284 (**Fig.6a**), a Wnt signaling activator and tubulin inhibitor [47], might form an LBHB with βGlu198. Crystal structure of tubulin-BML284 complex validated that BML284 binds to colchicine site and forms a hydrogen bond with βGlu198 (**Fig.6b and S7a**), which was proved to be an LBHB by QM/MM calculation as validated by a symmetric double well and low barrier potential energy curve and vibration energy splitting increase (ε _21 >_ ε _10_) (**Fig.6c and Table S2**). Superimposition of the tubulin–BML284 structure onto that of tubulin–PAC showed these two compounds overlap and both form an LBHB with βGlu198 in a similar manner (**Fig.S7b**). Immunoblotting results revealed BML284 indeed promotes tubulin heterodimers protein degradation in a concentration dependent manner while exhibits no effect on their mRNA levels (**Fig.6d and S7c**). The same as PAC, BML284-induced tubulin degradation could also be blocked by colchicine or PYR-41 (**Fig.6e and 6f**). TEM results confirmed that BML284 also induces spherical tubulin aggregates formation *in vitro* (**Fig.6g)**. Thiol Fluorescent Detection Kit Assay also indicated that BML284 induces purified tubulin denaturation (**Fig.6h**).Taken together, we identified a new tubulin degrader using the LBHB mediated degradation mechanism.

**Figure 6:**
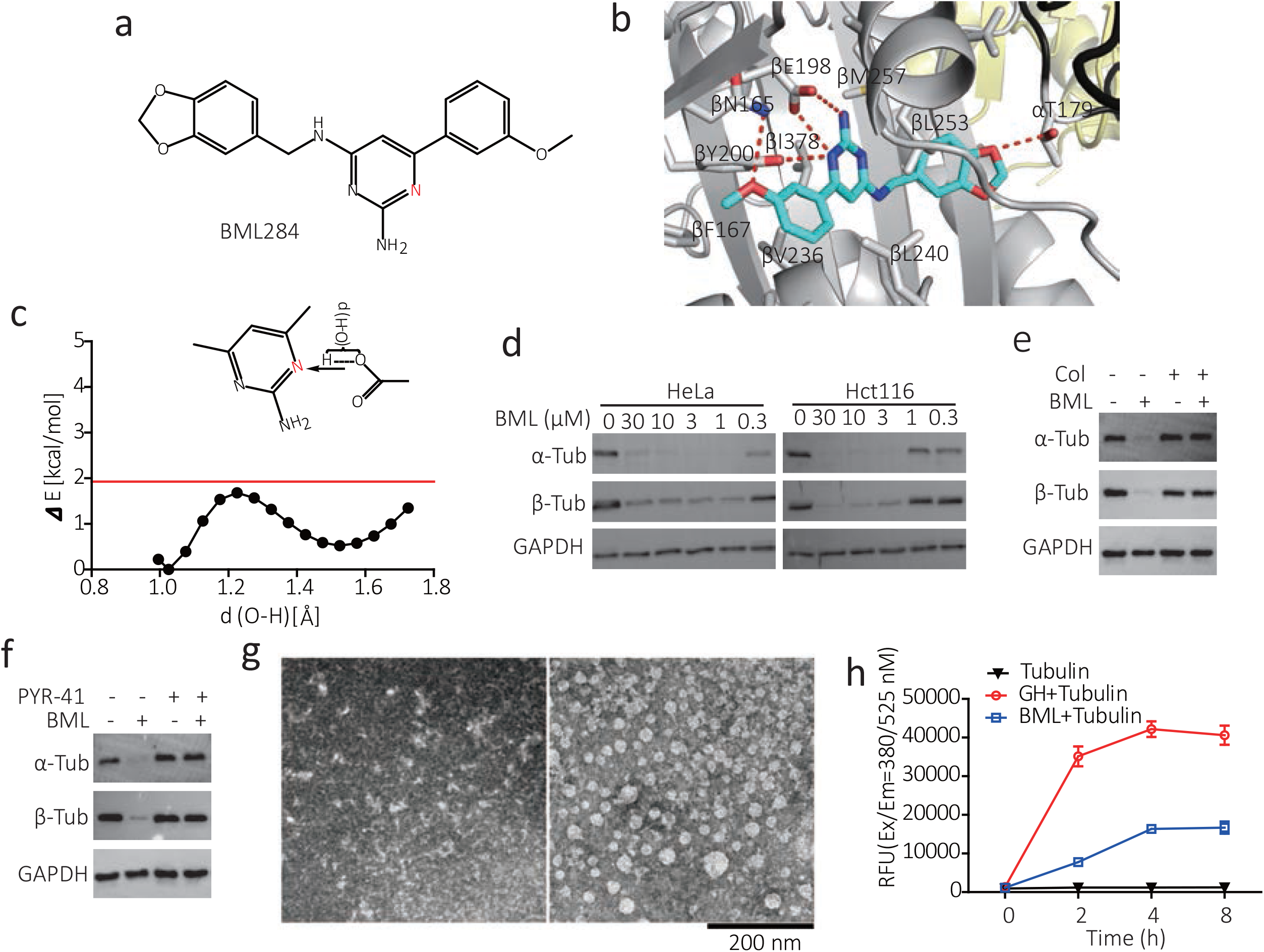
Idenfication of a new tubulin degrader with the LBHB mediated degradation mechanism. (**a**) Chemical structure of BML284. (**b**) Interactions between BML284 and tubulin. Residues that make interactions with BML284 are shown as scks and are labeled. Hydrogen bonds are drawn with red dashed lines. (**c**) Energy profiles along the proton transfer coordinate for H-bond donor-acceptor pairs of the indicated hydrogen bond in tubulin-BML284 complexes. The red line is the sum of zero-point vibraonal energy and kT (0.596 kcal/mol). (**d**) Immunoblot analysis of α- and β-tubulin protein levels in HeLa, Hct116 cells treated with the indicated concentrations of BML284 for 24 h. GAPDH was used as loading control. Results are representative of three independent experiments. (**e**) Colchicine blocks BML284-induced tubulin degradation. HeLa cells were pretreated with 3µM colchicine for one hour before treated with 3µM BML284 for 16 hours and then subjected to immunoblot analysis of α- and β-tubulin protein. Results are representative of three independent experiments. (**f**) Ubiquitin inhibitor PYR-41 rescue BML284-induced tubulin degradation. HeLa cells were pretreated 20 µM PYR-41 for one hour before treated with BML284 for 16 hours and then subjected to immunoblot analysis of α- and β-tubulin protein. Results are representative of three independent experiments. (**g**) Purified porcine tubulin (10 µM) was incubated with or without 20µM BML284 for 4 hours at room temperature, the images were taken by TEM. Results are representative of two independent experiments. (**h**) BML284 promotes purified tubulin denaturation. Tubulin (1µM) was incubated with guanidine hydrochloride(GH)or BML284 for 2, 4 or 8hours at room temperature. Then master reaction mix was added to the samples and incubated for another 30min. At last the fluorescent adduct were detected by a Biotech Gen5 spectrophotometer (1ex = 380/1em = 525 nm). Data are shown as the mean ± SD (n = 3).

## 3. Discussion

The human cell flags proteins for degradation by ubiquitinylation either through specific events (for example MDM2) [48] or as normal housekeeping to remove misfolded or denatured proteins [49]. Proteins with reduced stability are prone to unfolding and therefore more efficiently cleared in the normal housekeeping [50]. Molecules which promote protein ubiquitinylation will lead to proteolytic degradation of the targeted proteins are well established medicines [51].

Our data indicate there is a new method of specifically promoting target protein degradation. PAC, a known tubulin inhibitor [30], effectively destabilizes tubulin and thereby promotes its degradation as part of normal cell housekeeping. This effect is very specific since no widespread protein destabilization was observed and the effect is distinct from compounds such as isothiocyanates or withaferin A that also promote tubulin degradation. These compounds non-specifically bind to free protein thiols and promote degradation and / or alter the function of many proteins (isothiocyanates: Nrf, CDC25, GSH, MIF, STAT3 *et al* [52] and withaferin A: vimentin, actin HSP90, annexin II *et al* [53-55]), not just tubulin [56, 57].

Interestingly the co-complex of PAC with tubulin does not reveal any significant conformational changes that could easily explain the effect. However, structure-activity-relationship analysis using variants of PAC and site directed mutagenesis of β-tubulin, identifies a single key interaction, that between the pyridine nitrogen of PAC and βGlu198 (interior hydrophobic region of β-tubulin) as the trigger for degradation. Detailed calculations study show these two groups form an LBHB whilst other inhibitors, Noc and Pli, form only normal hydrogen bonds with βGlu198 and do not destabilize tubulin. We conclude that it is the LBHB between PAC and tubulin that destabilizes tubulin and consequently the protein is prone to unfolding and clearance via the house keeping ubiquitin system. The deep mechanism of LBHB mediate tubulin destabilization remains further investigation and here we proposed two possible hypothesizes: LBHB has unusual strong strength with some extent covalent character [46, 58], thus its abnormally strong strength could help to trap an unstable conformation of tubulin; Formation of an LBHB will release 10-20 kcal/mol energy and also the significant decrease in p*Ka* of βGlu198(from 9.6 to 2.3)will release about 10 kcal/mol energy as calculated by delta G° = 2.303RT(delta p*Ka*) [9, 59]. Therefore, a total 20-30 kcal/mol energy could make unfold of tubulin owing to the associated loss of net stability [60].

Using this LBHB mediated degradation mechanism we screened for compounds from lots of tubulin inhibitors with QM/MM calculation and molecular docking methods and found BML284, a Wnt signaling activator and tubulin inhibitor, can also form an LBHB with βGlu198 and promote tubulin degradation in a PAC-like manner as expected. Since LBHB was firstly proposed in the early 1990s, there have been few reports on other biochemical effects besides its roles in participating in enzyme catalysis and maintain protein structural integrity [10, 21, 22]. Our study confirms a new role of LBHB in mediating protein stability by small molecules, which could be utilized for novel drug discovery.

Molecules which form LBHB with buried residues represent a powerful new strategy in medicinal chemistry. These molecules are very different from the chaotropic agents that denature most proteins, they rely on specific protein ligand interactions. In another advantage, they are smaller and more “drug” like than the large proteolysis targeting chimaeras type molecules. The geometric constraints of the LBHB impose a high degree of selectivity upon those molecules thus reducing, at least in theory, unwanted side effects. As we have shown there exist excellent tools to judge whether any molecule participates in such an LBHB with a protein and measurement of the effect of such an LBHB upon protein stability is straightforward. These factors mean such an approach could be widely adopted.

## Acknowledgements

This work was founded by funds from National Natural Science Foundation of China (81872900, 81803021 and U1402222);1.3.5 project for disciplines of excellence, West China Hospital, Sichuan University; Post-Doctor Research Project, Sichuan University (2020SCU12036); China Postdoctoral Science Foundation Grants (2019T120855 and 2019M650248); Sichuan Science and Technology Program Grants (2019YJ0088, 2020YJ0089, 2019YFH0123 and 2019YFH0124).

## Author contributions

J.Y. performed most of the cellular and biochemical experiments and wrote the draft. Y.L. synthesized all chemical compounds. Q.Q. performed the tubulin mutation and animal experiments. R.W. and D.X. performed the computational calculations. W.Y, Y.Y, L.N., and Q.C. performed the structural biology experiments. H.P., H.W., L.O. and H.Y. analyzed the data. J.Y., Q.C., W.L. and L.C. conceived the idea and supervised the study. Q.C., and L.C. revised the manuscript.

All authors approved the final manuscript.

## Competing interests

The authors declare no competing financial interests.

## Correspondence and requests for materials

should be addressed to Q.C. or L.C.

## Data availability

Atomic coordinates and structure factors of tubulin complexed with PAC, Compound 9, Compound 11 or BML284 have been deposited in the Protein Data Bank under accession codes 7CDA, 7CE6, 7CE8 and 7CEK, respectively. All other data are available from the corresponding authors on reasonable request.

---

**Table S1.**
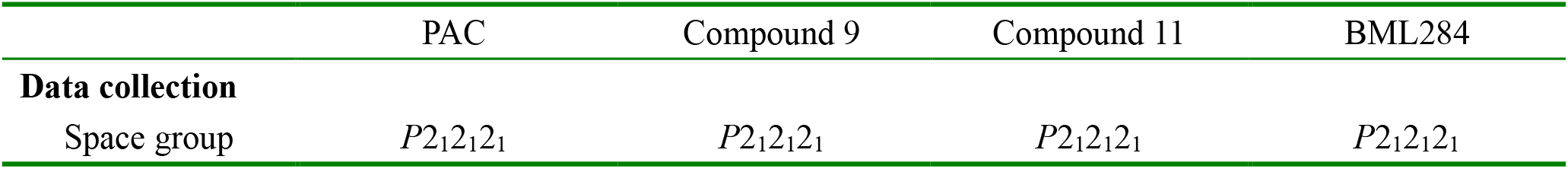
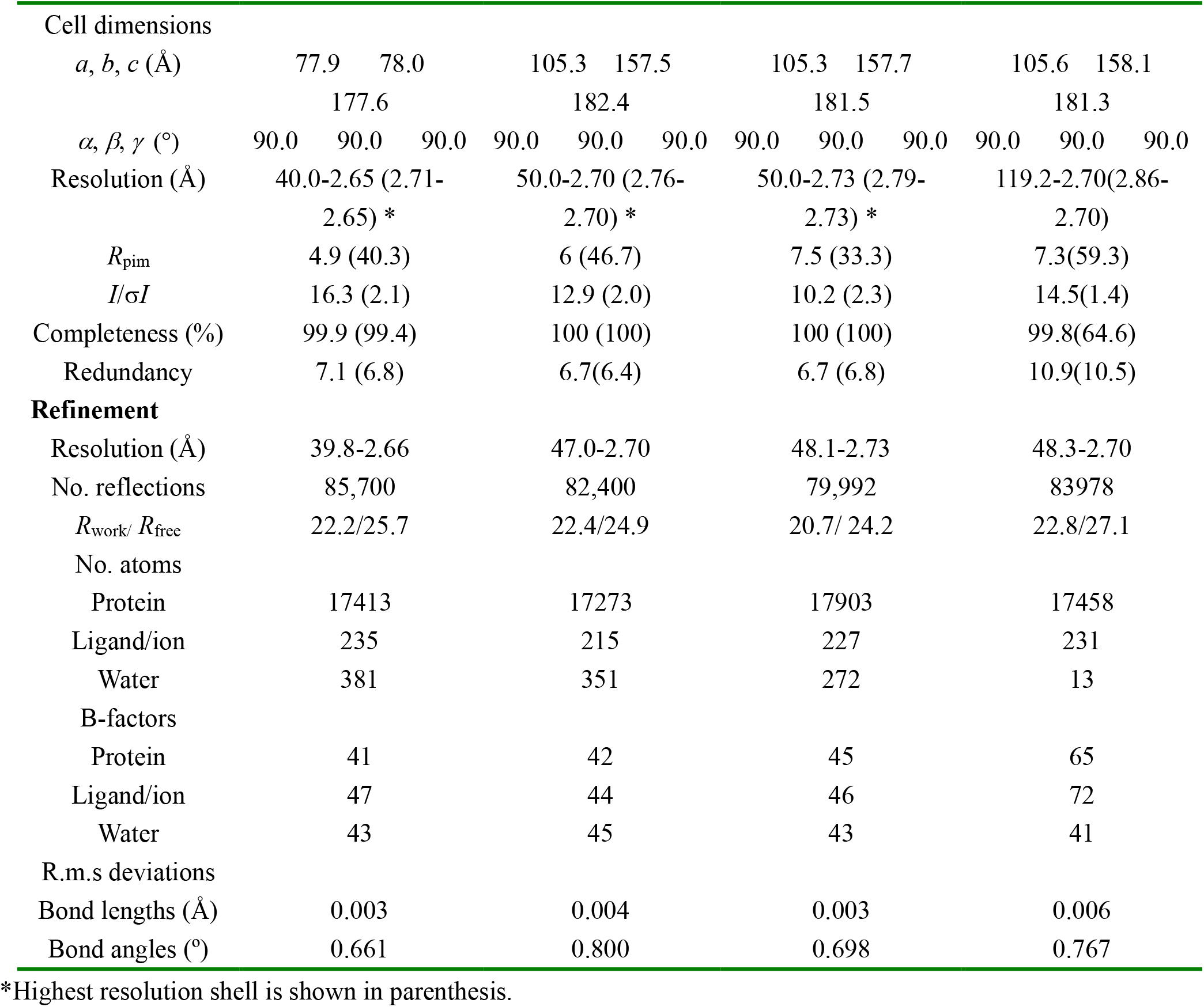
Data collection and refinement statistics.

**Table S2.** Vibrational energy levels (ε in kcal/mol) for the first three states of the proton in hydrogen bond between βGlu198 and compounds.

| | State | $\epsilon$ | $\Delta \epsilon$ | |
| --- | --- | --- | --- | --- |
| PAC | 0 | 1.46 |  |  |
| | 1 | 2.63 | $\epsilon_{10} = \epsilon_{\text{State 1}} - \epsilon_{\text{State 0}} = 1.17$ | $\epsilon_{10} < \epsilon_{21}$ |
| | 2 | 3.90 | $\epsilon_{21} = \epsilon_{\text{State 2}} - \epsilon_{\text{State 1}} = 1.27$ | |
| Noc | 0 | 1.87 |  |  |
| | 1 | 3.85 | $\epsilon_{10} = \epsilon_{\text{State 1}} - \epsilon_{\text{State 0}} = 1.98$ | $\epsilon_{10} > \epsilon_{21}$ |
| | 2 | 4.87 | $\epsilon_{21} = \epsilon_{\text{State 2}} - \epsilon_{\text{State 1}} = 1.02$ | |
| Pli | 0 | 3.20 |  |  |
| | 1 | 8.06 | $\epsilon_{10} = \epsilon_{\text{State 1}} - \epsilon_{\text{State 0}} = 4.86$ | $\epsilon_{10} > \epsilon_{21}$ |
| | 2 | 11.67 | $\epsilon_{21} = \epsilon_{\text{State 2}} - \epsilon_{\text{State 1}} = 3.61$ | |
| BML284 | 0 | 1.33 |  |  |
| | 1 | 2.34 | $\epsilon_{10} = \epsilon_{\text{State 1}} - \epsilon_{\text{State 0}} = 1.01$ | $\epsilon_{10} < \epsilon_{21}$ |
| | 2 | 3.82 | $\epsilon_{21} = \epsilon_{\text{State 2}} - \epsilon_{\text{State 1}} = 1.48$ | |

**Figure S1:**
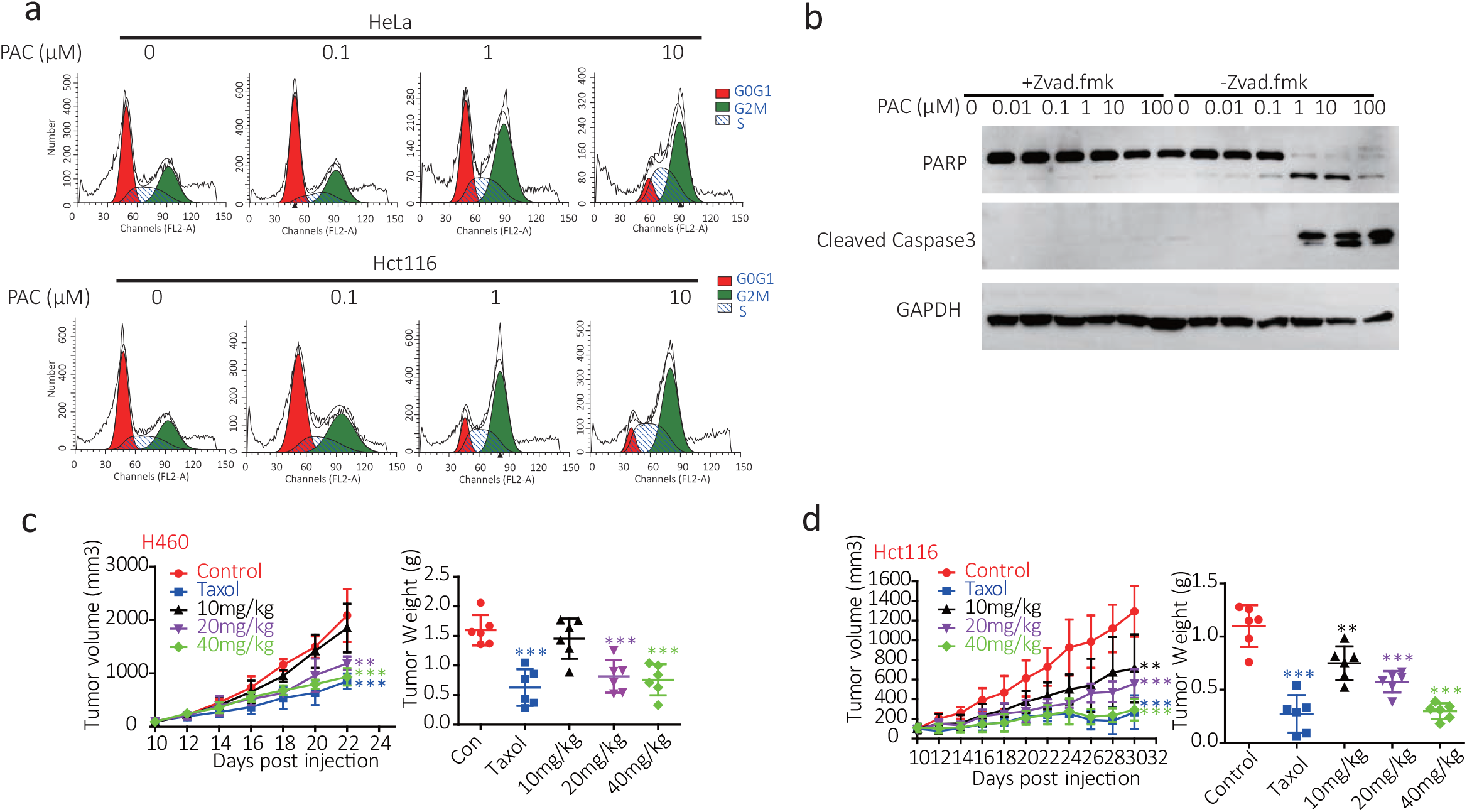
PAC induces G2/M phase cell cycle arrest and apoptosis and exhibits in vivo anticancer activity. (**a**) Flow cytometric histograms of various concentrations of PAC treated HeLa and Hct116 cells stained with propidium iodide. Results are representative of two independent experiments. (**b**) HeLa cells were treated with or without pan-caspase inhibitor Zvad.fmk for 1 hour before treated various concentrations of PAC for 48hous, then expression of poly ADP-ribose polymerase (PARP) and cleaved-caspase 3 were detected by immunoblot. GAPDH was loading control. Results are representative of two independent experiments. (**c**) In vivo antitumor activity of PAC (two days a time, tail vein injection) and taxol (30mg/kg, once a week, intraperitoneal injection) against H460 xenografts. Le: Time curve of tumor growth. Points showed as mean ± SD (n=6). Right: Bar chart of tumor weight. (**d**) In vivo antitumor activity of PAC (two days a time, tail vein injection) and taxol (30mg/kg, once a week, intraperitoneal injection) against Hct116 xenografts. Left: Time curve of tumor growth. Points showed as mean ± SD (n=6). Right: Bar chart of tumor weight.

**Figure S2:**
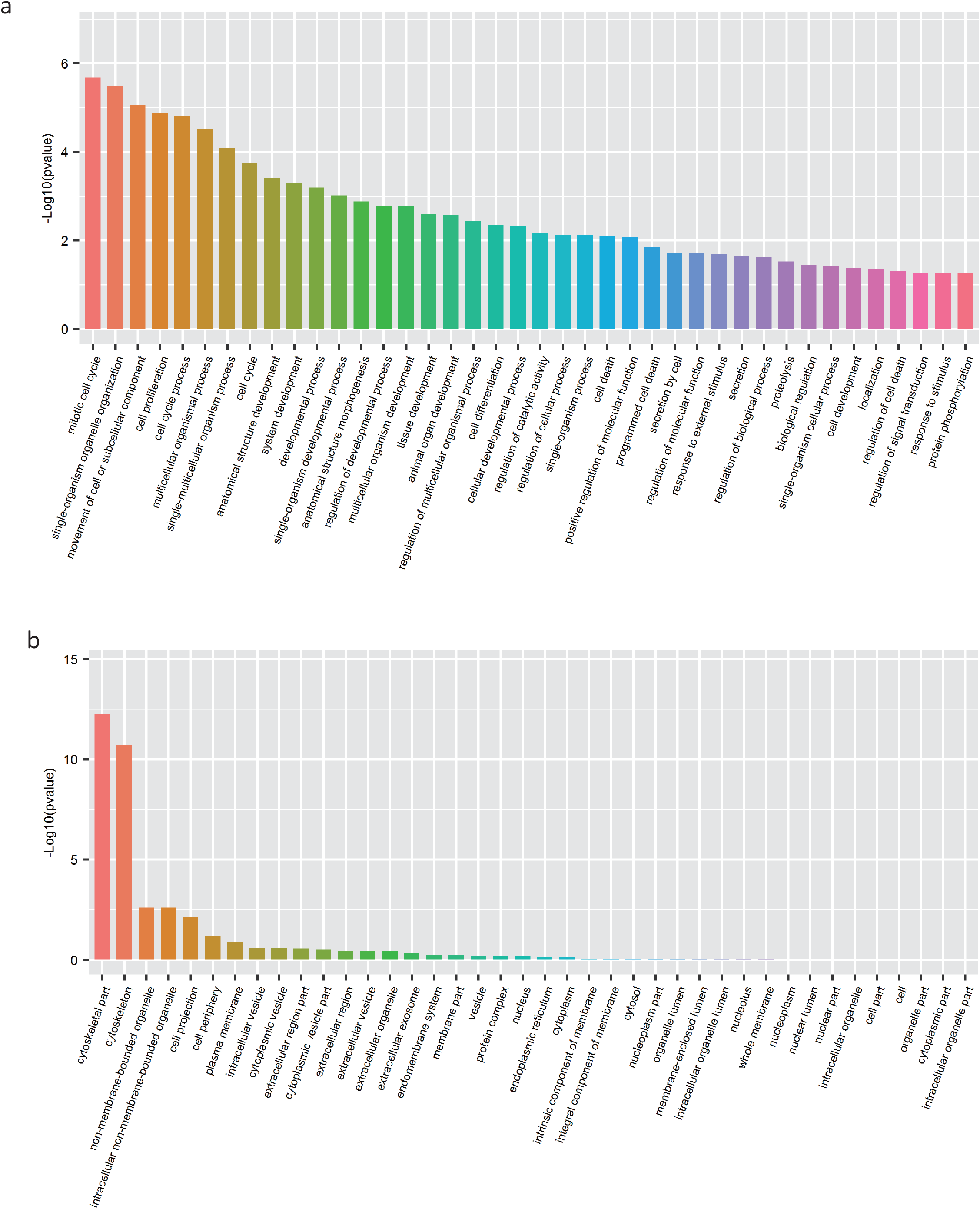
Biological processes and cellular components analysis of the quantitative proteomic data. (**a**) Rank order list of the most statistically enriched biological processes changes by PAC (Go Enrichment Analysis of biological processes) (**b**) Rank order list of the most statistically enriched cellular components changes by PAC (Go Enrichment Analysis of components changes).

**Figure S3:**
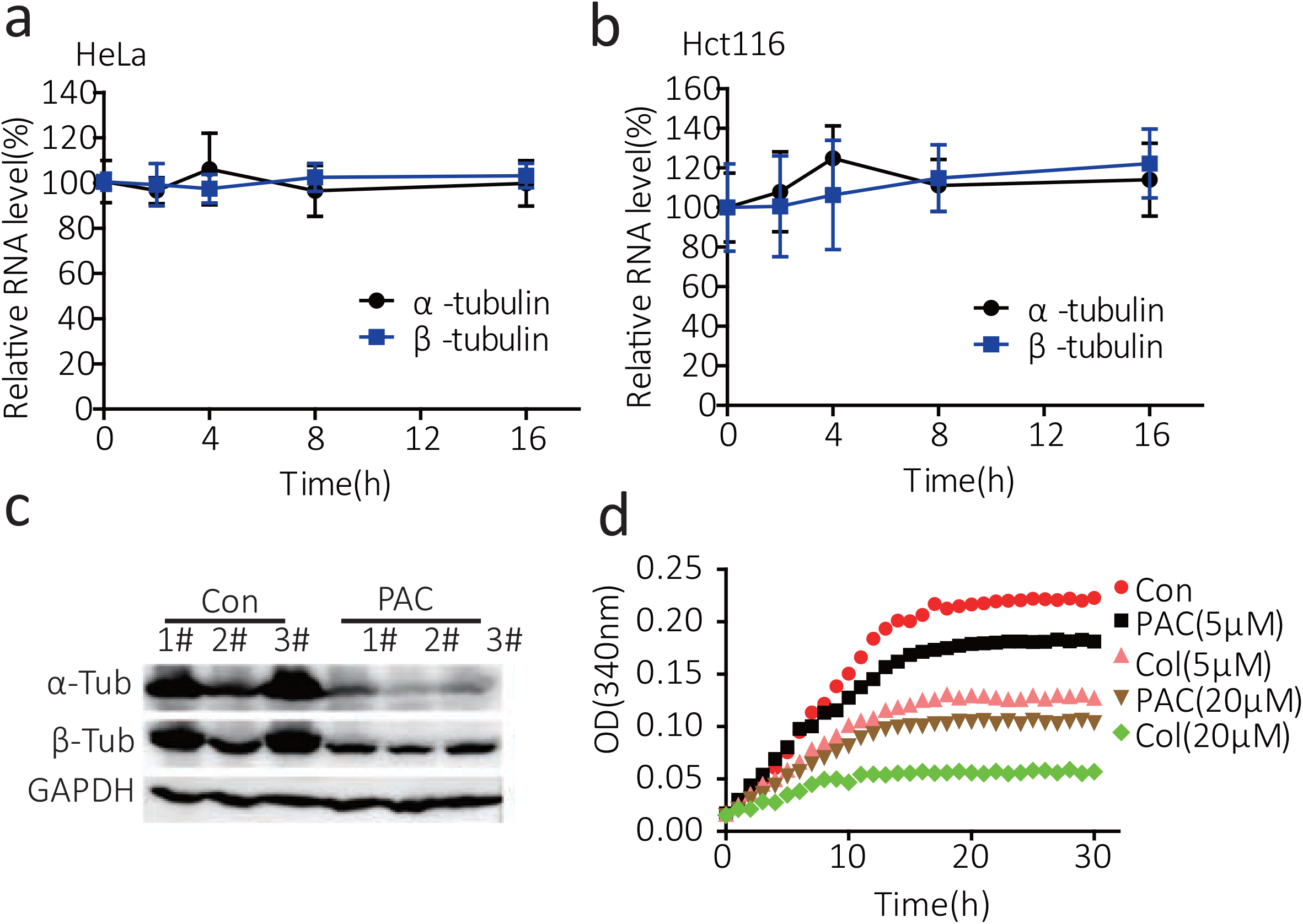
Supplementary figure for PAC induced tubulin degradation. (**a, b**) Time course of α- and β-tubulin mRNA levels in (**a**) HeLa and (**b**) Hct116 cells treated with 1 μM PAC or DMSO. Results are shown as the mean ± SD (n = 3). (**c**) Analysis of tubulin protein levels of Hct116 xenograft tissue. Tumor bearing mice were administrated with PAC at the dose of 80 mg/kg once through tail veil injection. Then mice were sacrificed after 24hours and xenograft tissues were collected for immunoblot to detect α- and β-tubulin protein. Results are representative of two independent experiments. (**d**) Indicated concentrations of compounds were co-incubated with tubulin (3mg/ml) at 37°C, absorbance at 340 nm were detected every one minutes for 30 minutes. Results are representative of two independent experiments.

**Figure S4:**
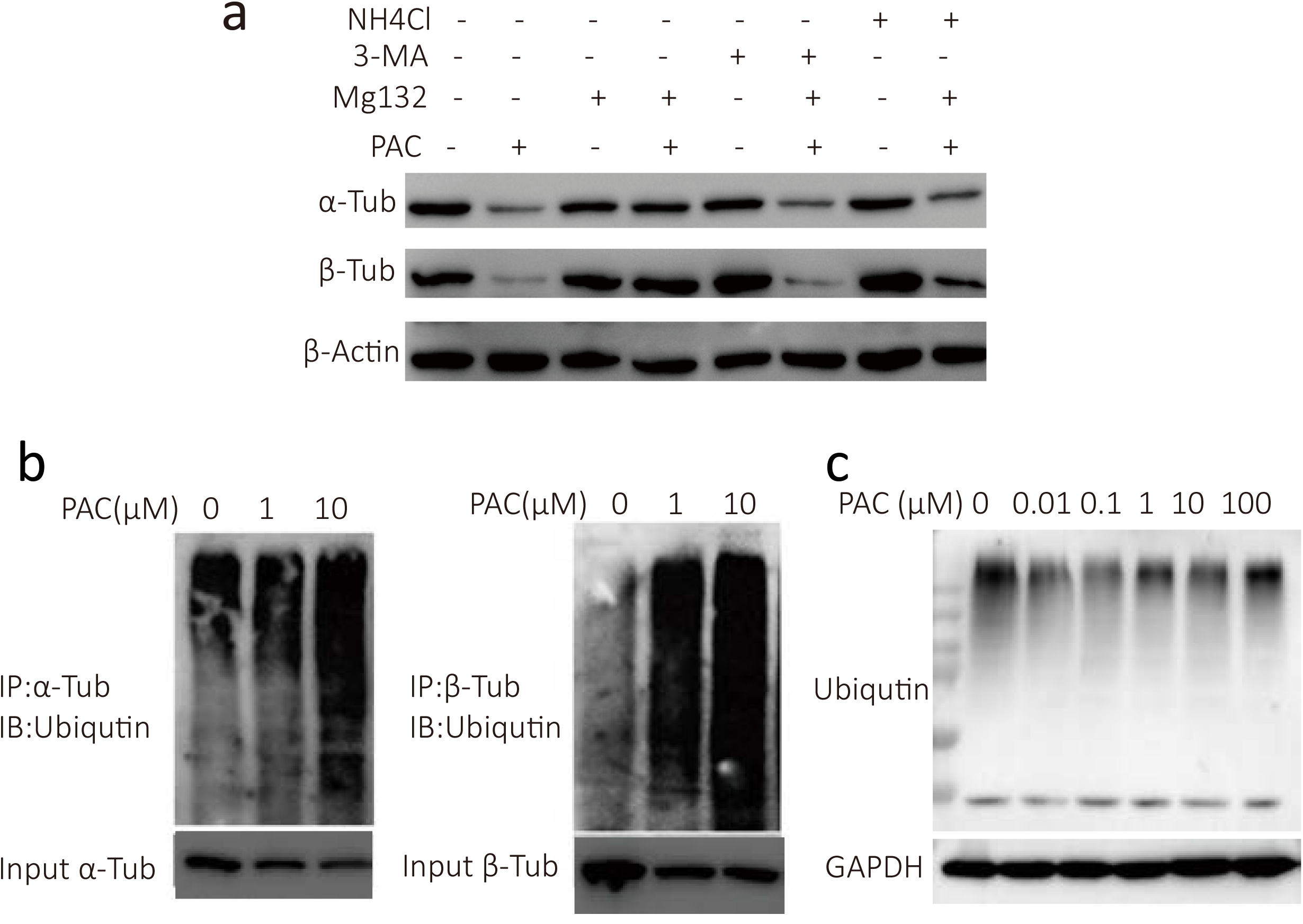
PAC promotes ubiquitin- and proteasome-dependent degradation of tubulins. (**a**) Chemical rescue of PAC induced tubulin degradation. HeLa cells were pretreated with Mg132, NH4Cl or 3-MA for 1hours before treated with PAC. Results are representative of three independent experiments. (**b**) HeLa cells were treated with 0, 1or 10μM PAC for 6hours, α- or β-tubulin were immunoprecipitated from the lysates, and ubiquitinylated α- or β-tubulin were detected with an-ubiquitin antibodies. Results are representative of three independent experiments. (**c**) HeLa cells were treated with indicated concentrations of PAC for 16hours, then total protein was extracted and analyzed by immunoblot for ubiquitin. Results are representative of two independent experiments.

**Figure S5:**
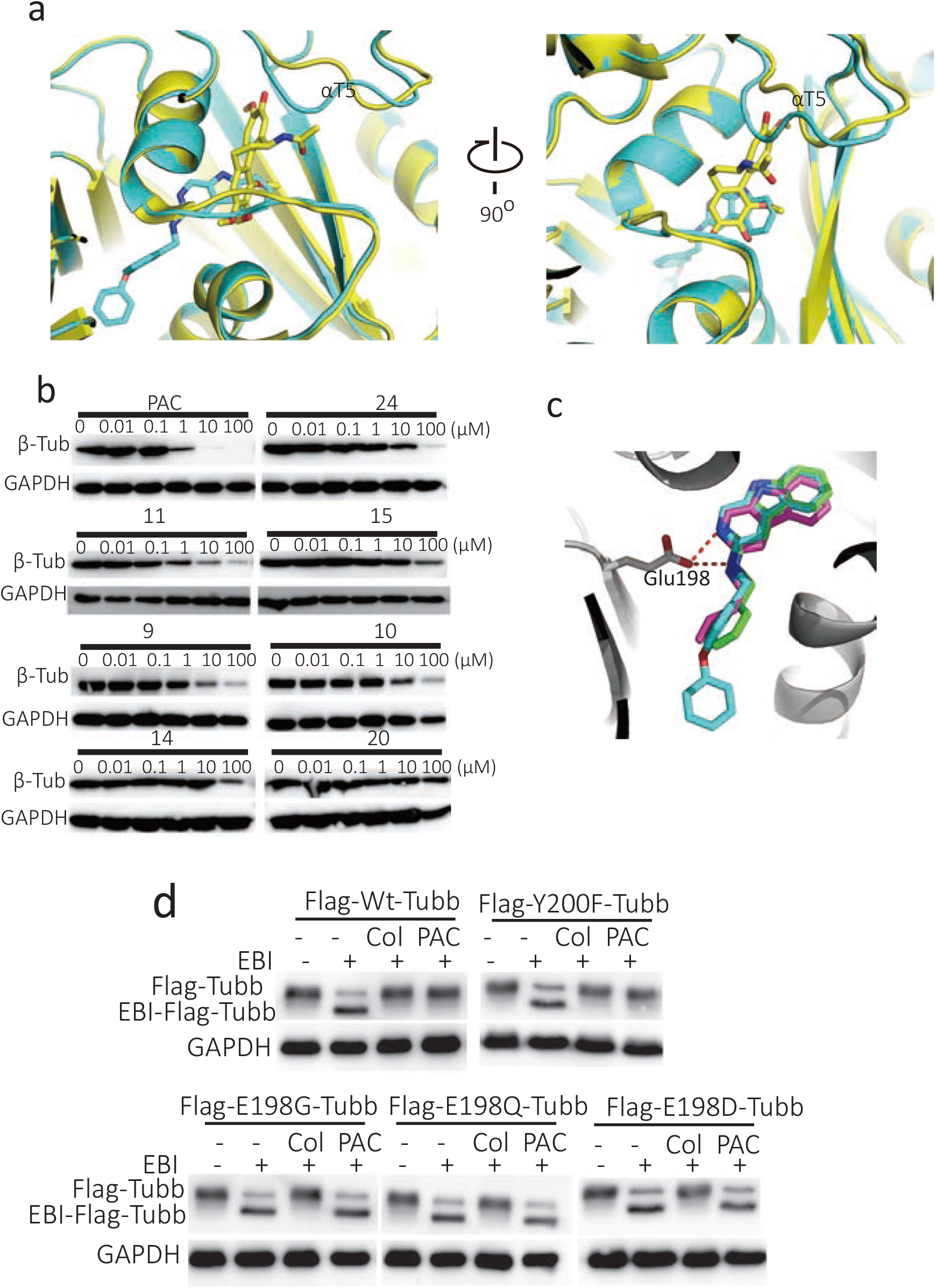
Supplementary figure to support an LBHB between PAC and βGlu198 mediates tubulin degradation. (**a**) Comparison of the tubulin binding modes between PAC and colchicine. The complex structures of tubulin-PAC (cyan) and Tubulin-colchicine (PDB ID: 4O2B, yellow) are superimposed. (**b**) Tubulin degradation activity of PAC derivatives. HeLa cells were treated with different concentrations of indicated compound for 16 hours, the β-tubulin protein levels were monitored by immunoblot. Results are representative of two independent experiments. (**c**) Comparison of the tubulin binding modes of PAC, compound 9 and 11. The structures of tubulin complexed with PAC (blue), compound 9 (green) and compound 11 (purple) are superimposed. (**d**) Identification the binding of PAC to wild type or mutant tubulins by EBI competition assay. MSCV-IRES-GFP vectors expressing both EGFP and wild type or mutant Flag-β-tubulin were transfected to HeLa cells for 24 hours. Then cells were treated with 10µM colchicine and 10µM PAC for 2hours before treated with 100µM EBI for another 2hours. Flag-β-tubulin level were detected by immunoblot. Results are representative of two independent experiments.

**Figure S6:**
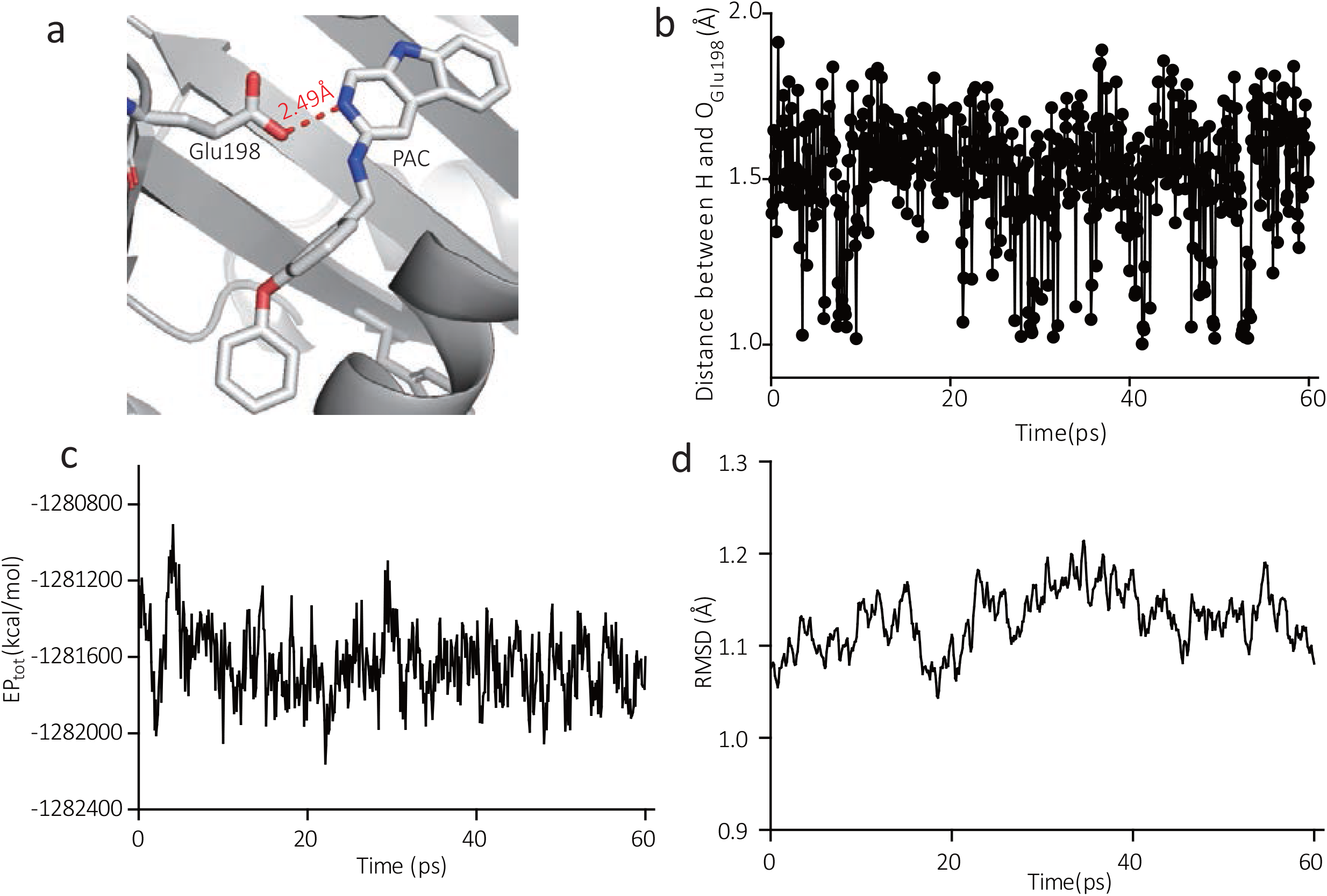
QM/MM MD simulation results. (**a**) Close-up view of the interaction between β Glu198 with PAC in QM/MM optimized structure showed the distance of the OGlu198-NPAC hydrogen bond. (**b**) The distance between proton and O of Glu198 in the 0 to 60 ps QM/MM MD simulation. (**c**) The total potential energy of the 0 to 60ps trajectory for PAC-tubulin complex. (**d**) Root mean square deviations (RMSDs) for 0 to 60 ps QM/MM MD simulation (QM: B3LYP/6-31G).

**Figure S7:**
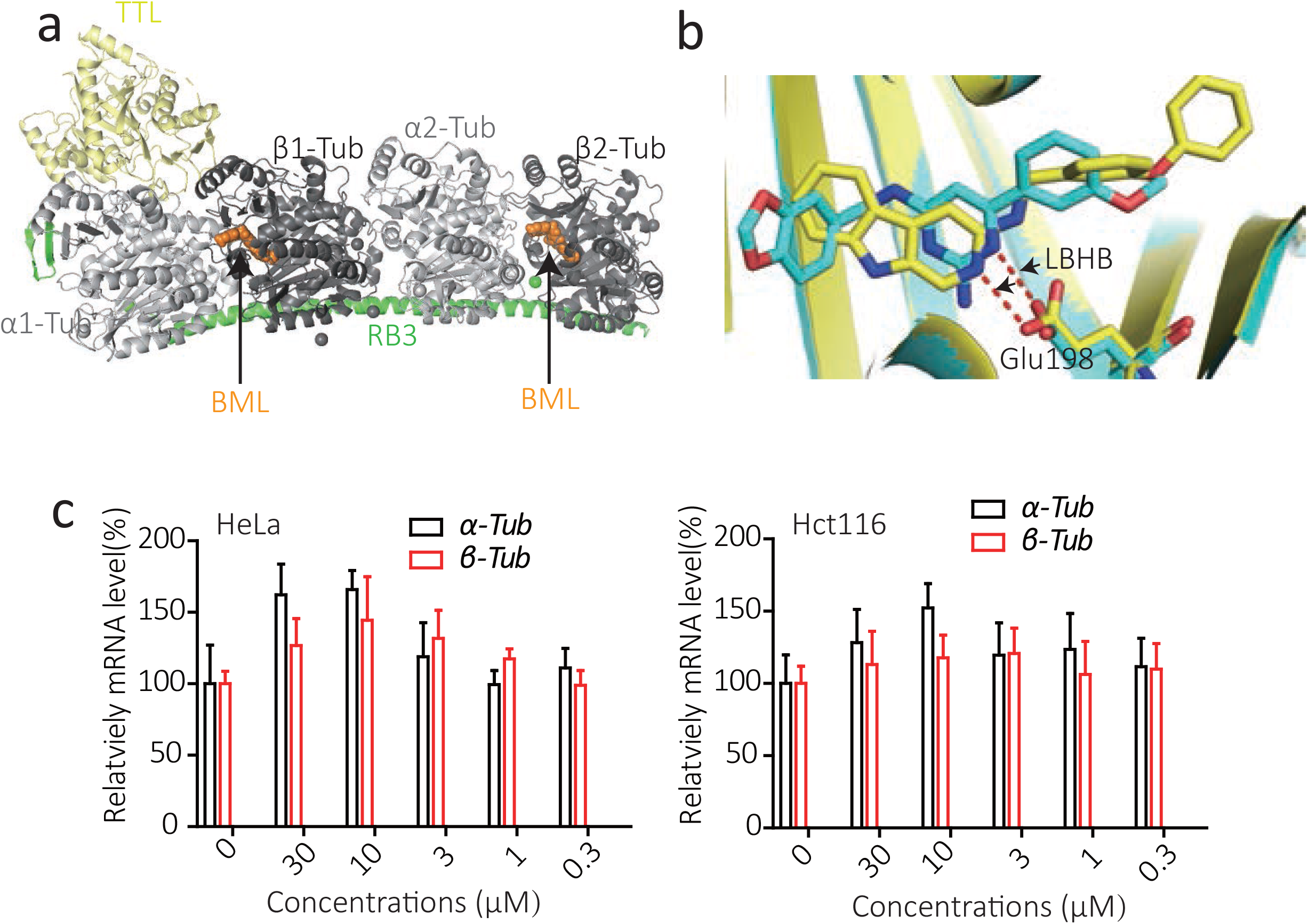
Supplementary figures to support BML284 induces tubulin degradation with the LBHB mediating degradation mechanism. (**a**) Overall structure of tubulin-BML284 crystal complex. The RB3-SLD is colored green, TTL is yellow, α-tubulin is grey, β-tubulin is black, BML284 is orange and shown as spheres. (**b**) Comparison of the tubulin binding modes of PAC and BML284. The structures of tubulin complexed with PAC (cyan) and BML284 (yellow) are superimposed. (**c**) The relative mRNA levels of α- and β-tubulin in HeLa and Hct116 cells treated with indicated concentrations of BML284 for 16hours were tested by Quantitative PCR, and the data are shown as the mean ± SD (n = 3).

## Notes

### Competing Interest Statement

The authors have declared no competing interest.

