## Supplemental Methods for "A new role of low barrier hydrogen bond in mediating protein stability by small molecules"

**Materials and Methods**

**1.Reagents**

Colchicine, paclitaxel, maytansine, vinblastine, β,γ-Methyleneadenosine 5′-triphosphate disodium salt (AMPPCP), DL-dithiothreitol (DTT), iodoacetamide (IAA), formic acid (FA), acetonitrile (ACN), methanol were obtained from Sigma；Pironetin were synthetized by Oskar Tropitzs； N,N'-ethylene-bis(iodoacetamide) (EBI) were obtained from Toronto Research Chemicals; Nocodazole，plinabulin，Mg132，PYR-41, 3-Methyladenine (3-MA) and Zvad.fmk were obtained from Selleck；Trypsin from bovine pancreas was purchased from Promega; Pierce BCA Protein Assay Kit from Thermo Scientific^TM^; Purified tubulin was purchased from Cytoskeleton, Inc.; Guanidine hydrochloride and other frequently used reagents were bought from Kelun Pharmaceutical.

**2.Chemistry**

TLC was performed on 0.20 mm silica gel 60 F_254_ plates (Qingdao Ocean Chemical Factory, Shandong, China). Visualization of spots on TLC plates was done by UV light. Melting points were recorded with a micro melting point tester (Neware Technology Ltd., Guangdong, China) and are uncorrected. NMR data were measured for ^1^H at 400 MHz and for ^13^C at 101 MHz on a Bruker Avance 400 spectrometer (Bruker Company, Germany) using TMS as an internal standard. High Resolution Mass Spectra (HRMS) were recorded on a Q-TOF Bruker Daltonics model IMPACT II massspectrometer (Micromass, Manchester, UK) in a positive mode.

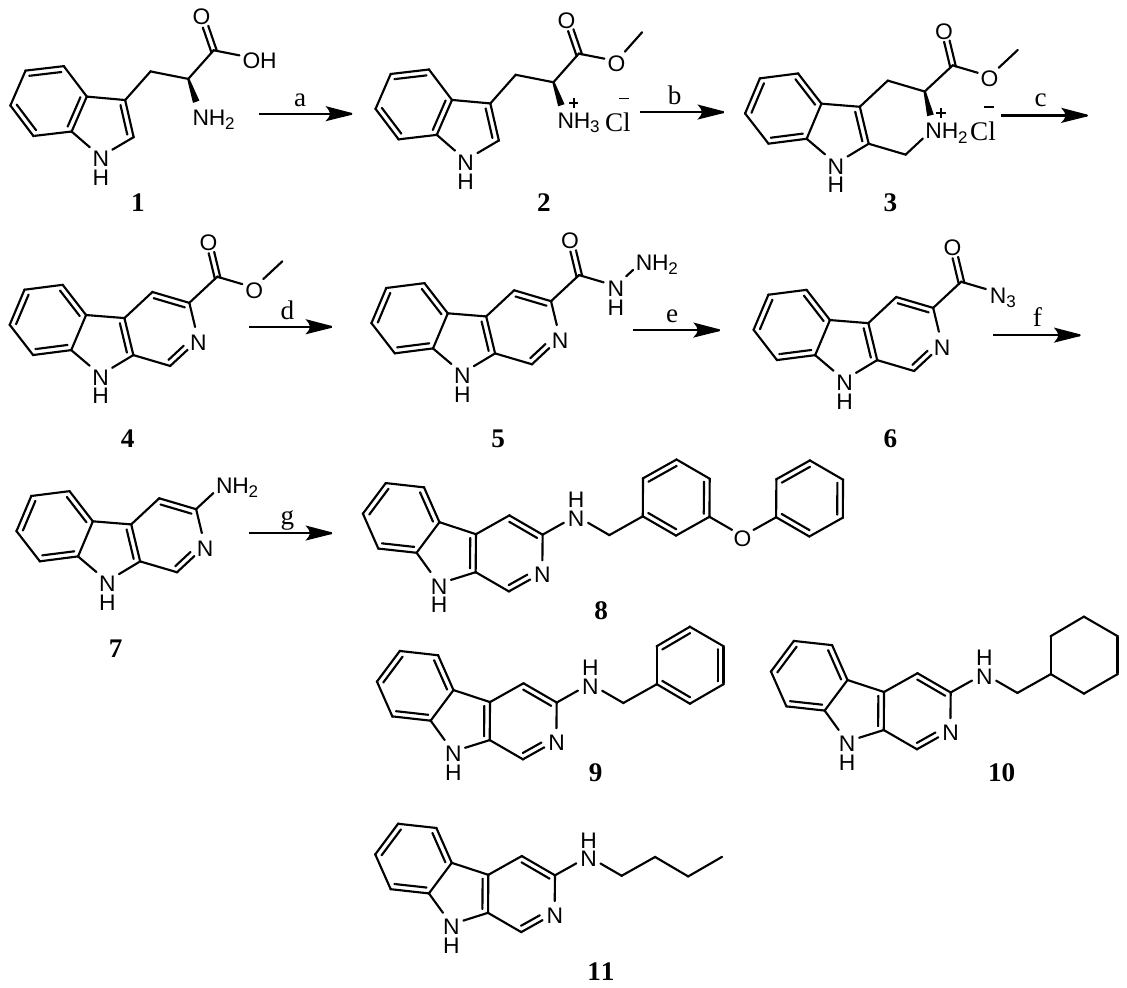
**Scheme 1.** Reagents and conditions: a) SOCl_2_, MeOH, 0-30℃ for 4h, then reflux; b) 37% HCHO, MeOH, reflux; c) TCCA, Et_3_N, DMF, -20 - 0 ℃; d) NH_2_NH_2_, MeOH, reflux; e) NaNO_2_, HCl, H_2_O, 0 ℃; f) HCl, H_2_O, reflux; g) 1) aldehyde, TFA, DMF, r.t.; 2) NaBH(OAc)_3_, MeOH, r.t.;

**2.1 General procedure for the synthesis of Methyl L-tryptophanate hydrochloride (2)**

To a suspension of 2.04 g (10 mmol) of commercial L-tryptophan (**1**) in 50 mL of MeOH was stirred at 0℃. Thionyl chloride (0.88 mL, 12 mmol) was added dropwise over 10 min. The reaction mixture was then allowed to warm to room temperature before being heated under reflux for 3h. The solvent and volatiles were evaporated under reduced pressure and the product was triturated with ethyl acetate to give the methyl ester hydrochloride salt as a colorless solid. The pure product was obtained in 95% yield. ^1^H NMR (400 MHz, DMSO-*d*_6_) δ 11.24 (s, 1H), 8.82 (s, 3H), 7.54 (d, *J* = 7.9 Hz, 1H), 7.39 (d, *J* = 8.1 Hz, 1H), 7.28 (d, *J* = 2.4 Hz, 1H), 7.13 – 7.05 (m, 1H), 7.04 – 6.94 (m, 1H), 4.19 (dd, *J* = 7.0, 5.4 Hz, 1H), 3.62 (s, 3H), 3.44 – 3.28 (m, 2H).

**2.2 General procedure for the synthesis of (S)-methyl 2,3,4,9-tetrahydro-1*H*-pyrido[3,4-b]indole-3-carboxylate hydrochloride (3)**

To a stirring solution of methyl L-tryptophen ester hydrochloride (24.5 g, 96.1 mmol) in methanol (250 mL) was added 37% aqueous formaldehyde solution (8.65g, 106.7 mmol) at room temperature. The resulting mixture was stirred at reflux for 3 h. The total volume was reduced in vacuo to approximately 50 mL, where upon a white precipitate formed. Methyl-tert-butyl ester (100 mL) was added, and the white suspension was stirred for 15 min. The resulting solid was received through filtration and then dried on high vacuum to afford pure compound **3** (22.3 g, 89%) as a fluffy white solid. ^1^H NMR (400 MHz, DMSO-*d*_6_) δ 11.25 (s, 1H), 7.48 (d, *J* = 7.8 Hz, 1H), 7.37 (d, *J* = 8.1 Hz, 1H), 7.15 – 7.06 (m, 1H), 7.05 – 6.96 (m, 1H), 4.63 (dd, *J* = 10.0, 5.3 Hz, 1H), 4.40 (s, 2H), 3.82 (s, 3H), 3.44 – 3.23 (m, 3H), 3.08 (dd, *J* = 16.0, 9.9 Hz, 1H).

**2.3 General procedure for the synthesis of methyl 9*H*-pyrido[3,4-b]indole-3-carboxylate (4)**

To a stirred solution of compound **3** (7.95 g, 34.5 mmol) and triethyl amine (9.09 g, 89.8 mmol) in 50 mL DMF, a DMF solution of trichloroiscyanuric acid (TCCA, 8.0 g, 34.5 mmol) was added slowly at -20℃. After the addition was completed, the reaction was allowed to warm slowly up to 0 ℃ with stirring for 2 h. The resulting product was precipitated from ice water, washed with ice water and dried in vacuo. The crude product yielding 80% as a gray solid and was used without further purification. ^1^H NMR (400 MHz, DMSO-*d*_6_) δ 12.06 (s, 1H), 8.97 (d, *J* = 0.9 Hz, 1H), 8.92 (s, 1H), 8.40 (d, *J* = 7.9 Hz, 1H), 7.67 (d, *J* = 8.2 Hz, 1H), 7.60 (ddd, *J* = 8.3, 6.9, 1.1 Hz, 1H), 7.32 (ddd, *J* = 8.0, 7.0, 1.1 Hz, 1H), 3.91 (s, 3H).

**2.4 General procedure for the synthesis of 9*H*-pyrido[3,4-b]indole-3-carbohydrazide (5)**

A solution of compound **4** (12.0 g, 50 mmol) in methanol (120 mL) containing hydrazine hydrate (85%, 20 mL) was refluxed for 6 h until there were no starting materials (TLC control). The resulting mixture was cooled, and the precipitate that formed was collected by filtration, washed with methanol (1 × 20 mL) and dried under vacuum to afford compound **5** (10.3 g, 85%). ^1^H NMR (400 MHz, DMSO-*d*_6_) δ 11.93 (s, 1H), 9.66 (s, 1H), 8.89 (d, *J* = 0.8 Hz, 1H), 8.83 (d, *J* = 0.8 Hz, 1H), 8.41 (d, *J* = 7.9 Hz, 1H), 7.66 (d, *J* = 8.2 Hz, 1H), 7.60 (ddd, *J* = 8.1, 6.8, 1.0 Hz, 1H), 7.30 (ddd, *J* = 8.0, 7.0, 1.1 Hz, 1H), 4.56 (s, 2H).

**2.5 General procedure for the synthesis of 9*H*-pyrido[3,4-b]indole-3-carbonyl azide (6)**

A suspension of the hydrazide **5** (12.1 g, 50 mol) in water (200 mL) was dissolved by the dropwise addition of concentrated HCl (10 mL). The pale yellow solution was cooled to 0 - 5 ℃ before the addition of a solution of sodium nitrite (3.6 g, 53 mmol) in water (10 mL). After stirring for 30 min at 0 ℃, the mixture was made basic with saturated aqueous NaHCO_3_, and the precipitate that formed was collected by filtration, washed with water, and dried in a dessicator under vacuum, yielding crude **6** (8.8 g, 75%) as a yellow solid with a tendency to decompose. The material was used without further purification for the following steps. ^1^H NMR (400 MHz, DMSO-*d*_6_) δ 12.27 (s, 1H), 8.99 (s, 1H), 8.97 (s, 1H), 8.41 (d, *J* = 7.9 Hz, 1H), 7.71 (d, *J* = 8.2 Hz, 1H), 7.68 – 7.61 (m, 1H), 7.37 (t, *J* = 7.4 Hz, 1H).

**2.6 General procedure for the synthesis of 9*H*-pyrido[3,4-b]indol-3-amine (7)**

A suspension of the azide **6** (11.85 g, 50 mmol) in a mixture of water (200 mL) and concentrated HCl (10 mL) was brought to reflux for about 1 h. When the starting material disappeared, the reaction mixture was cooled, the mixture was made basic with saturated aqueous NaOH. The precipitate that formed was collected by filtration and washed with water, and an analytical sample was recrystallized from EtOH, yielding **7** (6.4 g, 70%) as yellow powder. ^1^H NMR (400 MHz, DMSO-*d*_6_) δ 10.86 (s, 1H), 8.28 (s, 1H), 8.00 (d, *J* = 7.8 Hz, 1H), 7.42 (d, *J* = 6.1 Hz, 2H), 7.11 – 7.04 (m, 2H), 5.27 (s, 2H).

**2.7 General procedure for the synthesis of N-substitutedamino-β-carboline (8 - 11)**

Trifluoroacetic (TFA) acid (74 μL, 10.0 mmol) and corresponding aldehyde (1.1 mmol) were added to a solution of **7** (0.183 g, 1.0 mmol) in 5 mL dimethyl formamide (DMF) and the mixture was stirred for 2 h at room temperature. Then methanol (20 mL) was added to the reaction mixture, followed by NaBH(OAc)_3_ (1.06g, 5.0 mmol) and the mixture was stirred for an additional 5 h. The reaction was quenched by 6 N HCl (5 mL) and then cooled to room temperature, neutralized with saturated NaHCO_3_, and extracted with ethyl acetate (3 × 30 mL). The combined organic extract was washed with brine, dried with Na_2_SO_4_, and concentrated in vacuo. The residue was purified by silica gel chromatography to give pure **8** - **11**.

N-(3-phenoxybenzyl)-9*H*-pyrido[3,4-b]indol-3-amine (**8**)

yellow solid, yield: 71%. ^1^H NMR (400 MHz, DMSO-*d*_6_) δ 10.87 (s, 1H), 8.32 (s, 1H), 7.99 (d, *J* = 7.8 Hz, 1H), 7.42 (d, *J* = 6.1 Hz, 2H), 7.31 (t, *J* = 7.9 Hz, 3H), 7.18 (d, *J* = 7.6 Hz, 1H), 7.14 – 7.02 (m, 4H), 6.94 (d, *J* = 7.9 Hz, 2H), 6.86 – 6.78 (m, 1H), 6.49 (t, *J* = 6.3 Hz, 1H), 4.51 (d, *J* = 6.2 Hz, 2H). ^13^C NMR (101 MHz, DMSO-*d*_6_) δ 157.14, 156.94, 152.84, 144.41, 142.40, 131.68, 131.11, 130.38, 130.14, 128.41, 123.67, 122.85, 122.10, 120.94, 118.86, 118.46, 117.91, 116.99, 111.93, 96.15, 45.83. HRMS-ESI: calcd for C_24_H_20_N_3_O [M+H]^+^ 366.1607, found: 366.1608.

N-benzyl-9*H*-pyrido[3,4-b]indol-3-amine (**9**)

yellow solid, yield: 76%. ^1^H NMR (400 MHz, DMSO-*d*_6_) δ 10.85 (s, 1H), 8.33 (d, *J* = 0.9 Hz, 1H), 8.00 (d, *J* = 7.8 Hz, 1H), 7.46 – 7.36 (m, 4H), 7.29 (dd, *J* = 8.4, 6.9 Hz, 2H), 7.23 – 7.16 (m, 1H), 7.11 (s, 1H), 7.07 (ddd, *J* = 8.0, 6.2, 1.8 Hz, 1H), 6.44 (t, *J* = 6.3 Hz, 1H), 4.52 (d, *J* = 6.0 Hz, 2H). yellow solid, yield: 72%. M.P. °C. HRMS-ESI: calcd for C_18_H_16_N_3_ [M+H]^+^ 274.1345, found: 274.1346.

N-(cyclohexylmethyl)-9*H*-pyrido[3,4-b]indol-3-amine (**10**)

yellow solid, yield: 75%. ^1^H NMR (400 MHz, DMSO-*d*_6_) δ 10.79 (s, 1H), 8.32 (d, *J* = 1.0 Hz, 1H), 8.03 (d, *J* = 7.8 Hz, 1H), 7.45 – 7.37 (m, 2H), 7.11 – 7.03 (m, 2H), 5.77 (t, *J* = 5.7 Hz, 1H), 3.10 (t, *J* = 6.1 Hz, 2H), 1.86 – 1.77 (m, 2H), 1.74 – 1.66 (m, 2H), 1.64-1.55 (m, 2H), 1.32 – 1.12 (m, 5H). HRMS-ESI: calcd for C_18_H_22_N_3_ [M+H]^+^ 280.1814, found: 280.1810.

N-butyl-9*H*-pyrido[3,4-b]indol-3-amine (**11**)

yellow solid, yield: 54%. ^1^H NMR (400 MHz, DMSO-*d*_6_) δ 10.84 (s, 1H), 8.39 – 8.27 (m, 1H), 8.04 (d, *J* = 7.8 Hz, 1H), 7.50 – 7.34 (m, 2H), 7.15 – 6.98 (m, 2H), 5.72 (t, *J* = 5.7 Hz, 1H), 3.24 (q, *J* = 6.7 Hz, 2H), 1.58 (p, *J* = 7.3 Hz, 2H), 1.41 (dq, *J* = 14.4, 7.3 Hz, 2H), 0.93 (t, *J* = 7.3 Hz, 3H). HRMS-ESI: calcd for C_15_H_18_N_3_ [M+H]^+^, 240.1501, found: 240.1500.

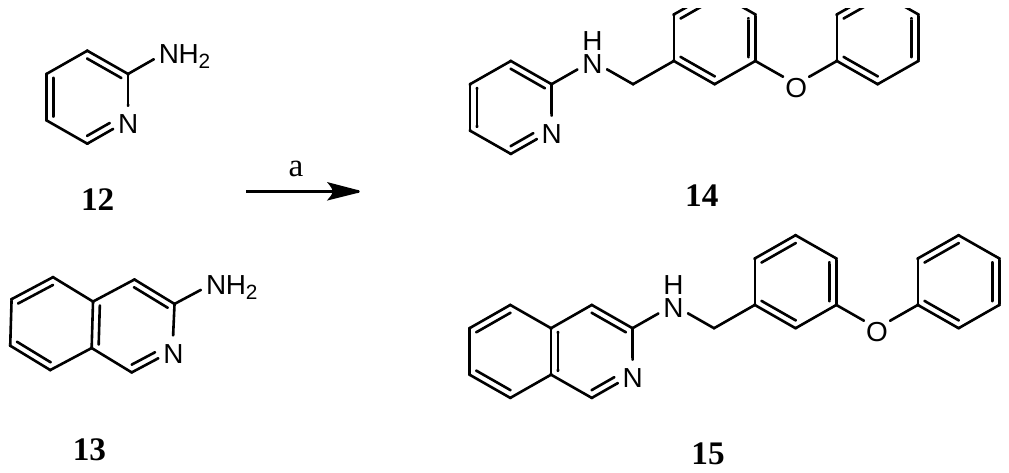

Scheme **2**. Reagents and conditions: a) 1) 3-phenoxybenzaldehyde, TFA, DMF, r.t.; 2) NaBH(OAc)_3_, MeOH, r.t.;

**2.8 General procedure for the synthesis of N-(3-phenoxybenzyl)pyridin-2-amine (14)**

The title compound **14** was synthesized in the same way as that of **8**. Yellow solid, yield: 93%. ^1^H NMR (400 MHz, DMSO-*d*_6_) δ 7.94 (dd, *J* = 5.1, 1.8 Hz, 1H), 7.40 – 7.26 (m, 4H), 7.12 (t, *J* = 7.2 Hz, 2H), 7.07 – 6.94 (m, 4H), 6.83 (dd, *J* = 8.0, 2.0 Hz, 1H), 6.53 – 6.43 (m, 2H), 4.47 (d, *J* = 6.1 Hz, 2H). HRMS-ESI: calcd for C_18_H_17_N_2_O [M+H]^+^ 277.1342, found: 277.1346.

**2.9 General procedure for the synthesis of N-(3-phenoxybenzyl)isoquinolin-3-amine (15)**

The title compound **15** was synthesized in the same way as that of **8**. Yellow solid, yield: 91%. ^1^H NMR (400 MHz, DMSO-*d*_6_) δ 8.84 (s, 1H), 7.79 (d, *J* = 8.1 Hz, 1H), 7.47 (dt, *J* = 14.8, 8.1 Hz, 2H), 7.31 (q, *J* = 7.5 Hz, 3H), 7.16 (t, *J* = 7.2 Hz, 2H), 7.07 (dd, *J* = 18.2, 9.3 Hz, 3H), 6.94 (d, *J* = 7.9 Hz, 2H), 6.84 (d, *J* = 7.7 Hz, 1H), 6.54 (s, 1H), 4.50 (d, *J* = 6.4 Hz, 2H). HRMS-ESI: calcd for C_22_H_19_N_2_O [M+H]^+^ 327.1498, found: 327.1496.

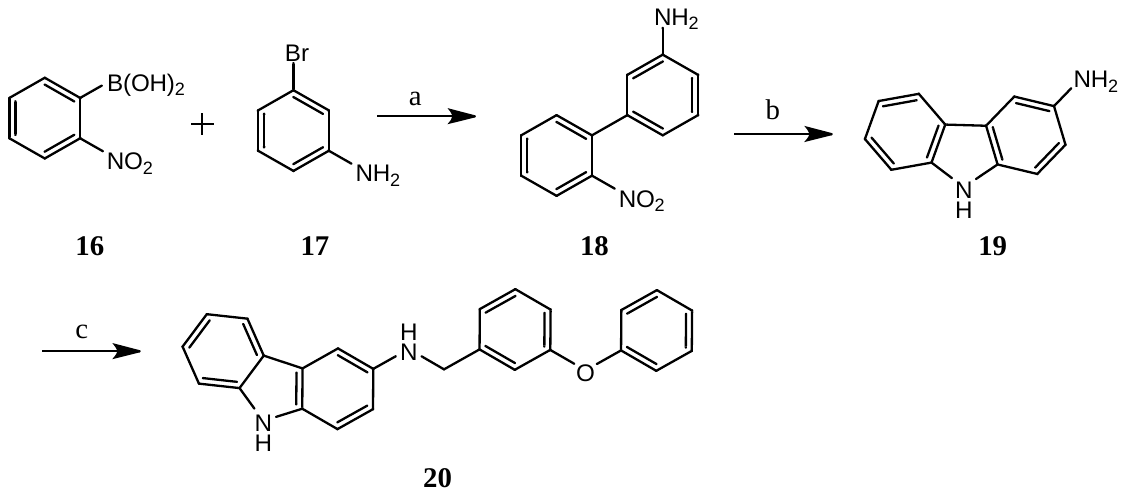

Scheme **3**. Reagents and conditions: a) Na_2_CO_3_, Pd(PPh_3_)_4_, 1,4-dioxane/H_2_O (4/1, v/v), 85 °C; b) PPh_3_, 1,2-dichlorobenzene, 180 °C; c) 1) 3-phenoxybenzaldehyde, TFA, DMF, r.t.; 2) NaBH(OAc)_3_, MeOH, r.t.;

**2.10 General procedure for the synthesis of 2'-nitro-[1,1'-biphenyl]-3-amine (18)**

Into a three-necked flask was added compound **16** (0.33 g, 2.0 mmol), compound **17** (0.38 g, 2.2 mmol), Na_2_CO_3_ (0.42 g, 4.0 mmol), Pd(PPh_3_)_4_ (0.12 g, 0.05 mmol) and 1,4-dioxane/H_2_O (4/1, v/v) (30 mL). The mixture was degassed by bubbling N_2_ for 15 mins, then the mixture was heated to 85 ℃for 8 h before quenched with water (250 mL). The resulting mixture was extracted with EtOAc (3 × 50 mL). The organic layer was washed by brine and dried with Na_2_SO_4_, filtered and evaporated under reduced pressure to afford a crude product, which was purified via a flash chromatography with silica gel to give compound **18** (0.31 g, 71% yield) as a yellow solid.

**2.11 General procedure for the synthesis of 9*H*-carbazol-3-amine (19)**

A solution of compound **18** (0.214 g, 1.0 mmol) and PPh_3_ (0.52 g, 2.0 mmol) in 1,2-dichlorobenzene (12 mL) was degassed by bubbling N_2_ for 15 mins, then heated to reflux with vigorous stirring for 24 h. The reaction was cooled to room temperature and concentrated under high vacuum. Chromatography of the yellow residue gave the product as a flaky, lustrous solid (0.15 g, 81% yield). ^1^H NMR (400 MHz, DMSO-*d*_6_) δ 10.68 (s, 1H), 7.89 (d, *J* = 7.8 Hz, 1H), 7.35 (d, *J* = 8.1 Hz, 1H), 7.27 (ddd, *J* = 8.2, 6.9, 1.1 Hz, 1H), 7.23 (d, *J* = 2.1 Hz, 1H), 7.19 (d, *J* = 8.5 Hz, 1H), 7.03 (ddd, *J* = 7.9, 7.0, 1.1 Hz, 1H), 6.76 (dd, *J* = 8.5, 2.2 Hz, 1H), 4.65 (s, 2H).

**2.12 General procedure for the synthesis of N-(3-phenoxybenzyl)-9*H*-carbazol-3-amine (20)**

The title compound **20** was synthesized in the same way as that of **8**. Yield: 92% yield, gray solid. ^1^H NMR (400 MHz, DMSO-*d*_6_) δ 10.72 (s, 1H), 7.87 (d, *J* = 7.7 Hz, 1H), 7.38 – 7.17 (m, 8H), 7.14 – 7.00 (m, 3H), 6.99 – 6.91 (m, 2H), 6.88 – 6.79 (m, 2H), 5.86 (t, *J* = 5.8 Hz, 1H), 4.35 (s, 2H).

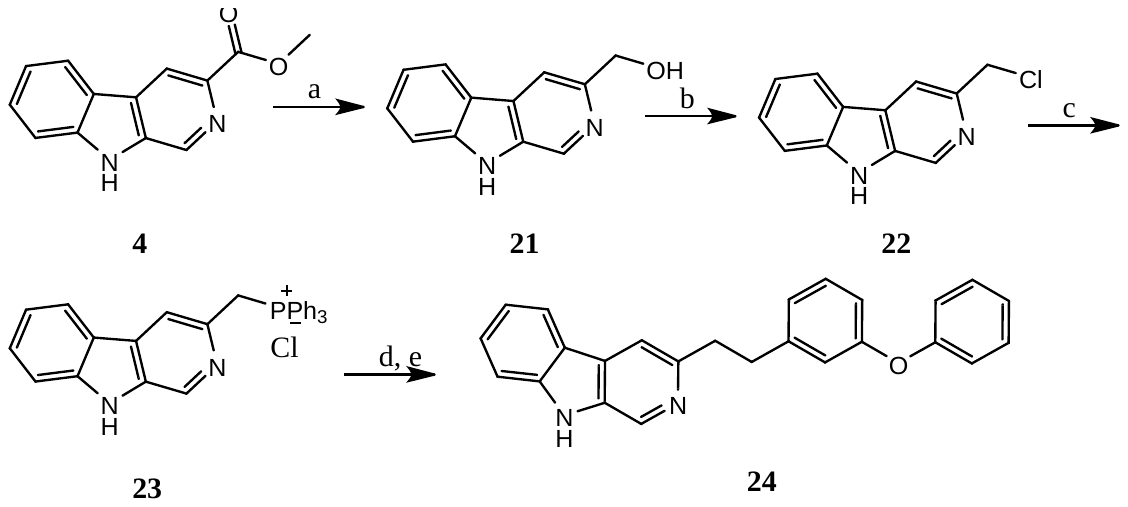
Scheme **4**. Reagents and conditions: a) LiAlH_4_, THF, 0 °C; b) SOCl_2_, reflux; c) PPh_3_, toluene, reflux; d) 3-phenoxybenzaldehyde, LiCl, LiOH, H_2_O, reflux; e) H_2_, Pd/C, EtOH, r.t.;

**2.13 General procedure for the synthesis of (9*H*-pyrido[3,4-b]indol-3-yl)methanol (21)**

To a solution of ester **4** (2.26 g, 10.0 mmol) in THF (100 mL) was added LiAlH_4_ (0.57 g, 15.0 mmol) in portions at 0 °C, then the mixture was stirred overnight at room temperature. After completion of the reaction, the mixture was quenched by water (10 mL) and stirred at room temperature for 2 h. Then the slurry was filtered, washed with dichloromethane, the filtrate was concentrated and purified via a flash chromatography with silica gel to afford **21** as a white solid (1.78 g, 90% yield). ^1^H NMR (400 MHz, DMSO-*d*_6_) δ 11.49 (s, 1H), 8.79 (d, *J* = 1.1 Hz, 1H), 8.24 (d, *J* = 7.8 Hz, 1H), 8.14 (s, 1H), 7.60 – 7.47 (m, 2H), 7.22 (ddd, *J* = 7.9, 6.8, 1.2 Hz, 1H), 5.33 (t, *J* = 5.8 Hz, 1H), 4.72 (d, *J* = 5.7 Hz, 2H). HRMS-ESI: calcd for C_12_H_11_N_2_O [M+H]^+^ 199.0872, found: 199.0870.

**2.14 General procedure for the synthesis of 3-(chloromethyl)-9*H*-pyrido[3,4-b]indole (22)**

3-(hydroxymethyl)-β-carboline (1.98g, 10.0 mmol) was added to an excess (20 mL) of thionyl chloride and the mixture was refluxed for 2 h. The excess thionyl chloride was then evaporated and the obtained residue was purified via a flash chromatography with silica gel to afford **22** as a white solid (1.99 g, 92% yield). ^1^H NMR (400 MHz, DMSO-*d*_6_) δ 11.77 (s, 1H), 8.89 (d, *J* = 0.9 Hz, 1H), 8.31 (s, 1H), 8.26 (d, *J* = 7.9 Hz, 1H), 7.62 (dt, *J* = 8.3, 1.0 Hz, 1H), 7.57 (ddd, *J* = 8.3, 6.9, 1.2 Hz, 1H), 7.27 (ddd, *J* = 8.0, 6.9, 1.2 Hz, 1H), 4.97 (s, 2H).

**2.15 General procedure for the synthesis of 9*H*-pyrido[3,4-b]indol-3-methyl phosphonium salt (23)**

Into a three-necked flask was added 3-(chloromethyl)-9H-pyrido[3,4-b]indole (0.433 g, 2.0 mmol), triphenylphosphine (0.525 g, 2.0 mmol) and toluene (20 mL). The mixture was degassed by bubbling N_2_ for 15 mins, then the mixture was refluxed for 6 h. The white solid formed was collected through filtration, washed with hot toluene and dried to give the desired product **23** that was used in the next steps without any further purification.

**2.16 General procedure for the synthesis of 3-(3-phenoxyphenethyl)-9*H*-pyrido[3,4-b]indole (24)**

To a suspension of phosphonium salts **23** (0.479 g, 1.0 mmol) in H_2_O (40 mL), 3-phenoxybenzaldehyde (0.297 g, 1.5 mmol), LiCl (0.42 g, 10.0 mmol) and lithium hydroxide hydrate (0.420 g, 10.0 mmol) were added. The mixture was refluxed for 6 h and then extracted with EtOAc (3 x 25 mL). The organic layer was washed with brine and dried over Na_2_SO_4_ and concentrated under reduced pressure. The crude product was purified by a flash chromatography on silica gel to give mixtures of E and Z stilbenes with ratios approximatively 70 : 30. These mixtures were further reduced by H_2_ in EtOH with the catalyst Pd/C. After completion of reaction, the mixture was filtered and the filtrate was concentrated under reduced pressure. The obtained residue was purified via a flash chromatography with silica gel to afford **24** as a white solid. (0.258 g, 71% yield over two steps). ^1^H NMR (400 MHz, DMSO-*d*_6_) δ 11.45 (s, 1H), 8.80 (d, *J* = 1.0 Hz, 1H), 8.15 (d, *J* = 7.9 Hz, 1H), 7.88 (s, 1H), 7.57 (dt, *J* = 8.3, 1.1 Hz, 1H), 7.52 (ddd, *J* = 8.2, 6.8, 1.2 Hz, 1H), 7.31 – 7.18 (m, 4H), 7.11 – 7.01 (m, 2H), 6.92 – 6.84 (m, 2H), 6.82 – 6.76 (m, 2H), 3.22 – 3.11 (m, 2H), 3.11 – 3.02 (m, 2H). HRMS-ESI: calcd for C_25_H_21_N_2_O [M+H]^+^ 365.1655, found: 365.1654.

**3 Antibodies.**

The following antibodies were used: Anti-α tubulin antibody（Abcam，ab52866，for immunoblot，1:1000 dilution or ZENBIO， #250009，for immunoblot，1:5000 dilution），Anti-β tubulin antibody（Abcam，ab6046，for immunoblot，1:1000 dilution or ZENBIO， #200806， for immunoblot，1:5000 dilution），Anti-GAPDH antibody（Abcam，ab8245，for immunoblot，1:5000 dilution） Anti-Flag antibody (Proteintech, 20543-1-AP, for immunoblot, 1：1000), Anti-GFP antibody（Cell Signaling Technology, #2555, for immunoblot，1:1000 dilution），Anti-Cleaved caspase 3 antibody(ZENBIO，#341034，for immunoblot，1:2000 dilution)，anti-PARP antibody （Santa Cruz Biotechnology，sc-74470, for immunoblot，1:200 dilution）, anti-ubiquitin antibody（Santa Cruz Biotechnology, sc-47721, for immunoblot，1:200 dilution）.

**4 TMT Quantitative Proteomics**

**4.1 Sample Preparation**

Hela cells were seeded in six-well plates for 24 hours and then treated with or without 1μM PAC for six hours，then cells were collected and lysed with RIPA buffer (containing 2mM PMSF and proteinase inhibitor mixture). Then samples were centrifuged at 10,000 × g for 5 minutes to pellet cell debris. Supernatant were collected and preserved for further analysis. Protein concentrations of the supernatant were determined by the BCA Protein Assay Kit. We have done three biological repetition.

**4.2 Trypsin digesting and Labeling with the TMT ® Reagents – 10plex**

The samples were reduced and blocked by cysteine before digested with trypsin solution at 37 °C overnight. Allow each required vial of TMT Reagent–10plex (ThermoFisher Scientific, USA) to reach room temperature. Spin to bring the solution to the bottom of the vial. Add 41μL of anhydrous ACN to each room-temperature vial. Vortex each vial to mix, then spin and dissolved for 5 min. Then the 20μL TMT solution was added to each vial and incubated at room temperature for 60min. 4 μL of 5% hydroxylamine was added to each vial to stop the reaction for 15min. Then the three control samples were labeled with TMT^10^-126, TMT^10^-127N and TMT^10^-127C, respectively. The three PAC treated samples were labeled as TMT^10^-128C, TMT^10^-128N and TMT^10^-129C, respectively. At last, the contents of each labeled sample were combined into one vial.

**4.3 Nano LC-MS/MS Analysis**

A Dionex Ultimate 3000 Nano LC system coupled with an Orbitrap Fusion™ Tribrid™ Mass Spectrometer (Thermo Fisher Scientific, USA) with an ESI nanospray source was used. The spin columns (Thermo Fisher Scientific, USA) were conditioned twice with 0.1% TFA solution. Then digested samples were fractionated with elution solutions (20%ACN,80%TFA solution~30%ACN, 70%TFA solution~40%ACN, 60%TFA solution~50%ACN, 50% TFA solution). The liquid contents of each samples were evaporated to dryness using vacuum centrifugation. Each fractionated sample was loaded to a nanocolumn (100 μm×10 cm in-house made column packed with a reversed-phase ReproSil-Pur C18-AQ resin (3 μm, 120 Å, Dr. Maisch GmbH, Germany) on a nanoflow UPLC (Easy-nLC1000 (ThermoFisher Scientific, USA)). The loaded sample volume was 5 μL and the flow rate was set at 600 nL/min. The mobile phase was composed of solvent A(0.1% formic acid in water) and solvent B which changed follow times: from 6% to 9% B for 15 min, from 9% to 14% B for 20 min, from 14% to 30% B for 60 min, from 30% to 40% B for 15 min and from 40% to 95% B for 3 min, eluting with 95% B for 7 min. After purification by UPLC, peptides were then subjected to Orbitrap Fusion™ Tribrid™ Mass Spectrometer (Thermo Fisher Scientific, USA) for analysis. Up to top 15 most intense peptide ions from the preview scan in the Orbitrap.

**5.Cell culture**

Hela, Hct116, H460 and SU-DHL-6 Cells were obtained from the KeyGEN Biotech Co. (Nanjing, Jiangsu, China). Hela, Hct116, and SU-DHL-6 cells were cultured in DMEM (containing 10% fetal bovine serum and 1% penicillin-streptomycin) and H460 cells were cultured in RPMI-1640 (containing 10% fetal bovine serum and 1% penicillin-streptomycin). Cells were incubated in a humidified incubator in 37°C with 5% CO_2_. All cell lines were authenticated by STR testing and was free of mycoplasma.

**6. Cell viability detection**

Cells were seed on 96-well plates and cultured for 24hours and then cells were treated with different compounds at various concentrations for 48 hours, then 20 μL MTT reagents (100μg/ml) were added to each well and incubated for 2-3hours. Culture medium were removed and 150 μL DMSO were added to each sample for 10min on a rotator. At last, the OD (optical density) values were measured at 570 nm on a microplate reader (Biotek, USA ).

**7. Cell Cycle analysis**

Cells were seeded on six-well plates for 24hours before treated with different concentrations of PAC for 16hours. Cells were collected and washed with PBS for two times, then fixed with 75%(v/v) pre-cold ethanol for 12hous at 4°C. Cells were washed three times with PBS and then stained with PI staining buffer (PI, 50 mg/ml) for 30 min. Then, samples were subjected to cytometry for cell cycle analysis.

8. **Vector construction**

The Flag tag was fused to the C-terminus of TUBB（β-Tubulin）, TUBB(Y200F) and TUBB(E198G) genes, and PacI and BamHI restriction sites were added at all ends. The complete sequence was synthesized by Genewie (Suzhou, China) and cloned into MSCV-IRES-GFP expression vector. Mutations (E198Q and E198D) of TUBB were performed using the Q5 Site-Directed Mutagenesis kit (NEB #E0554S). The specific primers used for point mutation is: TUBB(E198Q), Mutation primer, AGAGAATACTGATCAGACCTATTGCATTG, Reverse primer: ACCAACTGATGGACGGAGAGG; TUBB(E198D) Mutation primer, AGAGAATACTGATGACACCTATTGCATTG, Reverse primer: ACCAACTGATGGACGGAGAGG.

8. **Transfection**

Hela cells were seed on six well plates and cultured for 24hous before being transfected by Lipofectamine 2000 reagent. Approximately 250μL OptiMEM (Thermo) containing 2.0μg plasmid DNA, and 250μL OptiMEM containing 7.5μL Lipofectamine 2000 reagent were prepared and incubated for 5min，respectively， before mixed together and incubated for another 20min. Then the mixture was added to Hela cells and cultured for 24hours before adding the drugs for 16hours. Then total proteins were extracted and analyzed by western blot using flag and GFP antibodies, GAPDH was employed as loading control.

**10.Western Blot**

**10.1 Sample preparation**

For *in vitro* study, tumor cells were seeded on six-well plate and cultured for 24hours, then treated with different compounds for different time. All cells were washed with PBS (1 min) and then digested with trypsin，and then collected and washed with PBS (1 min) again. Then 2🞨loading buffer (diluted from 6🞨 loading buffer by RIPA lysis buffer) are added to cells for lysis for 10min, and the samples were then boiled for 10 min.

For *in vivo* study, Hct116 cells (2🞨10^6^) were seeded to the right flank of six BALB/c mice. After the tumor volume reached about 250mm^3^, mice were randomly divided into two groups (control group and PAC group) with 3 mice each. Mice in PAC group were treated with 80mg/kg PAC (dissolved in physiological saline containing 5% ethanol and 5% tween-80) through tail vein injection, and the control group were treated with equal volume of physiological saline containing 5% ethanol and 5% tween-80. After 24 hours, mice were sacrificed and tumors were excised immediately and frozen in liquid nitrogen. The tumor tissue in liquid nitrogen was crushed and transferred to an EP tube containing RIPA lysate (containing PMSF and other protease inhibitors) for ultrasonication for 5 min and then continued to lyse on ice for 30 min. After centrifugation at 4°C(13000rpm) for 30 min, the supernatant were mixed with loading buffer and then boiled for 10 min.

**10.2 Western detection**

Equal amount of samples were loaded to SDS-PAGE for electrophoresis, then protein on SDS-PAGE were transferred to PVDF (polyvinylidene difluoride,0.22μm) membranes electrophoretically at 4°C for 2 hours. Then the PVDF membranes were blocked by 5% skim milk (diluted in 1🞨loading buffer) for 1 hour, and the first-antibodies (diluted in blocking buffer ) were incubated with membranes for 12hours at 4°C. Membranes were washed with 1🞨PBST (3🞨10min) to remove the unbounded antibodies, and incubated with second-antibodies for 45min. Membranes were washed with 1🞨PBST (3🞨10min) again to remove unbounded second-antibodies. At last, membranes were incubated with enhanced chemiluminescence reagents (Millipore) and imaged using a fully automatic chemiluminescence image analysis system (Tianneng, China).

**11 Immunofluorescence**

Cells were seeded on microscope cover glass placed in six-well plate for 24hours before treated with different compounds for pre-set times, then cover glasses were washed with PBS for 2min before incubated with 50% methanol/50% acetone for 3min. Cells were incubated with fist-antibody (diluted in 3% BSA buffer) for 4hours at room temperature, then washed with PBS(4🞨10min) and incubated with Alexa Fluor-conjugated second antibody and 10% DAPI for 1hour, then washed again with PBS (4🞨10min). At last, cover glass was mounted onto microscope slides with a small drop of glycerol. The samples were imaged using a fluorescence microscope (Olympus, Japan).

**12. Quantitative-PCR**

Cells were seed on six-well plates for 24hours culture and then treated with PAC for 2, 4, 8 or 16hours, total mRNA were extracted with TRIzol (Invitrogen) agents according to the manufacturer’s protocol. The quality of total mRNA were determined by a NanoDrop1000 spectrophotometer (Thermo Fisher Scientific). High Capacity cDNA Reverse Transcription Kit (Applied Biosystems) were used for the cDNA synthesis following the manufacturer’s instructions. Quantitative PCR analysis were performed using the Taq Universal SYBR Green Supermix (BIO-RAD) on an CFX96 Real-time PCR System (BIO-RAD). CFX Manager software version 2.1 (BIO-RAD) were used to determine the cycle threshold (Ct) values. PCR efficiency and relative quantitation were calculated by a six-point standard curve. Relative expression of *α-tubulin* and *β-tubulin* were normalized against the expression level of *β-actin*. The primers are list below :

*α-tubulin*：Forward Primer:TCGATATTGAGCGTCCAACCT;

Reverse Primer:CAAAGGCACGTTTGGCATACA;

*β-tubulin*：Forward Primer：TGGACTCTGTTCGCTCAGGT;

Reverse Primer：TGCCTCCTTCCGTACCACAT；

*β-actin*：Forward Primer: CATGTACGTTGCTATCCAGGC;

Reverse Primer: CTCCTTAATGTCACGCACGAT.

**13. Gel filtration assays**

Purified tubulin were dissolved to 3 μM in PEM buffer (80 mM PIPES, pH 6.9, 0.5 mM EGTA, 2 mM MgCl_2_) (Containing 1mM GTP)，before incubated with different compounds for different times. After incubation, samples were analyzed by a gel filtration assay. This assay was performed with a Pharmacia Akta chromatography system, using a Superdex 200 10/300 GL gel fltration column (GE Healthcare, Piscataway, NJ), with a 200μL injection loop. The column flow rate was 0.5 ml/min. Running buffer: 20 mM Pipes, 200 mM KCl, 1 mM MgCl_2_, 50 mM GTP (Sigma, St. Louis, MO), 0.1% β-Mercaptoethanol (v/v), adjust pH to 6.5 using KOH. Filter the running buffer using a 0.22-μm filter. Injection volumue:100μL. Detection wavelength: 280nm.

**14. Tubulin polymerization assay**

This assay was performed according to previous reported study[1]. Briefly, porcine tubulin (3mg/ml) were dissolved in PEM buffer (80 mM PIPES, pH 6.9, 0.5 mM EGTA, 2 mM MgCl_2_) with 15% glycerol and pre-treated with different compounds at different concentrations or vehicle DMSO on 4°C，then all samples were added 1mM GTP and transferred to microplate reader (Biotek, USA ) to detect the OD value at 340 nm at 37 °C every minute for 30min.

**15. EBI competition assay**

N,N'-ethylene-bis(iodoacetamide) (EBI), a homobifunctional thioalkylating agent, was used to crosslink the Cys-239 and the Cys-354 residues of β-tubulin involved in the colchicine-binding site[2]. The covalent binding of EBI to β-tubulin forms an adduct that is easily detected by western blotting as a second immunoreactive band of β-tubulin that migrates faster than the native β-tubulin band on SDS-PAGE[2]. Cells were treated with MG132 for 1hour before incubated with or without different concentrations of compounds for 2 hours and then incubated with 100 μM EBI for another 2hours，then total proteins were extracted and incubated with loading buffer before denatured in boiling water for 10min. β-Tubulin band and EBI-β-tubulin band were detected by western blot using β-tubulin antibody.

**16.Transmission electron microscopy**

Purified porcine tubulin (10 μM) were dissolved in PEM buffer at 4℃，then pre-treated with or without colchicine（20μM）for 30min before incubated with or without 20μM PAC for 4hours at 4°C. Then 5μL samples were added to carbon films supported 300-mesh per inch formvar for 2 min, then washed three times with water before negatively stained for 30 s with 2% (w/v) phosphotungstic acid. Then images of tubulin on formvar were observed on a transmission electron microscope (A FEI T12，80Kv), and a Serial EM software was employed to take pictures.

**17.Fluorometric thiol quantitation assay**

Thiol Fluorescent Detection Kit (Enzolifesciences, ADI-907-036) provides an accurate method to quantify free thiol groups in proteins. The proprietary thiol sensor generates a strongly fluorescent adduct (λex = 380/λem = 525 nm) upon reacting with a thiol-containing compound. To detect whether PAC and other compounds induced denaturation of tubulin and BSA, we tried to detect whether compounds induced thiol exposure by this assay according to the manufacturer’s protocol. Accordingly, Tubulin (1μM) was incubated with guanidine hydrochloride（positive control）or other compounds or DMSO for 2, 4 or 8hours at 4℃. Then Master Reaction Mix were added to the samples and incubated for another 30min. At last the fluorescent adduct were detected by a microplate reader (Biotek, USA ).

**18. Tubulin stability determination by Differential Scanning Fluorimetry（DSF）.**

Tubulin stability was tested using nanoDSF (Prometheus NT.48). Purified tubulin（0.2mg/ml）in PEM buffer was incubated with different compounds for 20 mins. Then capillaries were directly immersed into the protein solution to load the sample for testing. The heating temperature range is set at 20-95 °C with 1℃/min increase. Tryptophan fluorescence at 330 and 350 nm (emission wavelengths) were detected (330F and 350F). Melting temperatures (Tm value) were obtained by calculating the maximum of the first derivative of the F350/F330 fluorescence ratios.

**19. Structural Biology**

**19.1 Protein expression and purification.**

The complex of two tubulins, one stathmin-like domain of RB3 (RB3–SLD) and one tubulin tyrosine ligase (TTL) (the T2R–TTL complex) was produced as described with slight modifications [3]. RB3–SLD was overexpressed in *Escherichia coli* BL21(DE3), purified sequentially by anion-exchange chromatography (QFF; GE Healthcare) and gel filtration (Superdex 75, GE-Healthcare). The purified protein was concentrated to 10 mg/mL and stored at 80 °C until use. TTL was overexpressed in E. coli BL21(DE3) and purified by nickel-affinity chromatography (BeaverBeads™, Beaverbio) followed by gel filtration (Superdex 200, GE-Healthcare). Purified TTL in Bis-Tris propane (pH 6.5), 200 mM NaCl, 2.5 mM MgCl2, 5 mM β-mercaptoethanol and 1% glycerol was concentrated to 20 mg/mL and stored at 80 °C until use. Porcine brain tubulin (Cytoskeleton, Catalog # T-238P) was supplied at 10 mg/mL (buffer: 80 mM Pipes, pH 6.9, 2.0 mM MgCl2, 0.5 mM EGTA and 1 mM GTP) and stored at 80 °C until use. The T2R–TTL complex was prepared by mixing tubulin, RB3–SLD and TTL in a 2: 1.3:1.2 molar ratio, and then 1 mM β,γ-methyleneadenosine 50-triphosphate disodium salt, 5 mM tyrosine and 10 mM DTT were added and the complex was concentrated to 20 mg/mL at 4 °C.

**19.2 Crystallization and crystal soaking.**

The T2R–TTL crystals were obtained at 20 °C in a buffer consisting of 6% PEG4000, 8% glycerol, 0.1 M MES (pH 6.7), 30 mM CaCl_2_ and 30 mM MgCl_2_. Seeding method was used to obtain single crystals. Rod-like crystals appeared after 2 days and grew to maximum dimensions within 1 week. For crystal soaking, 0.1 μL of small-molecule inhibitor (dissolved in DMSO at 10 mM concentration) was added to a 2 μL crystal-containing drop for 12 h at 20 °C.

**19.3 Data collection and structure determination.**

The reservoir solution supplemented with 20% (v/v) glycerol was used as the cryoprotectant. The crystals were transferred into the cryoprotectant for a few seconds, and then mounted in nylon loops and flash-cooled in liquid nitrogen. Diffraction data were collected on beamline BL19U1 of National Facility for Protein Science Shanghai (NFPS) at Shanghai Synchrotron Radiation Facility (Shanghai, China). Data were processed using HKL3000 [4]. The structures were determined by molecular replacement method using the T2R–TTL structure (PDB ID: 4I55) as a search model. The refinement was performed using COOT [5] and PHENIX [6]. The model quality was checked with MOLPROBITY [7].

**20.** **Computational details**

Low barrier hydrogen bonds (LBHBs) have been proposed to play roles in protein functions during enzymatic catalysis [8, 9]. Tubulin-PAC (PDB:7CDA), tubulin-plinabulin (PDB:5C8Y), tubulin-nocodazole (PDB:5CA1) complexes, tubulin-BML284 (PDB:7CEK) were used here for LBHB caculation. Of course, in order to quantitatively understand and confirm the LBHB occurring in this special protein, high level quantum mechanics computations are required to shed some insights into this issue.

**20.1 QM/MM Optimization**

To tackle bond formation and dissociation process occurred in the protein active site, combined quantum mechanics and molecular mechanics (QM/MM) method represents one of the powerful tools in this field.

First, the geometry optimization for all complexes were carried out using our own n-layered integrated molecular orbital molecular mechanics (ONIOM) method [10-12]. A two-layered ONIOM method implemented in Gaussian09 suite of program[13] was employed in the geometry optimization and subsequent electronic potential energy scanning calculations. Details of ONIOM can be found elsewhere[10]. Basically, the total energy of the system can be described as summation of three terms as following:

$E^{ONIOM}=E^{High, model}+E^{Low, real}-E^{Low,model}$ , (1)

in which “High” and “Low” represent the levels of computational method. “Real” denotes the whole system, which is calculated using the low-level method. “Model” contains the part that needs to be treated using both the high and low levels. To include solvent effect, the whole system was then solvated in a pre-equilibrated rectangular water box with sodium ions added to neutralize the system. The total number of atoms for the final system is calculated to be around 26000. Density functional theory (DFT) with B3LYP exchange correlation function was employed in this work for the geometric optimization. During the geometric optimization, a standard basis set of 6-311G (d, p) was employed. A universal force field (UFF) [14] was applied to account for all atoms belonging to low layer including protein environment and water molecules.

**20.2 DFT Models**

To confirm the existence of LBHB in the systems we investigated, the truncated active site model with high level of quantum chemistry methods should be enough. For Tubulin-PAC (PDB: 7CDA), tubulin-plinabulin (PDB: 5C8Y), tubulin- nocodazole (PDB:5CA1) and tubulin-BML284 (PDB: 7CEK) complexes, the truncated models include Glu198 neutral charged side chain (protonated) and Tyr200. The reaction coordinate for LBHB reaction can be defined as the distance between the Oε atom of Glu198 and hydrogen atom. A series of structures were optimized for one-dimensional potential energy curves for the migration of the hydrogen nucleus along the reaction coordinate. For DFT computation, ωB97X-D/6-311++G** level [15, 16] of theory was employed in this work. One-dimensional potential energy curves for the migration of the hydrogen nucleus along the direction vector from Glu198 to ligand have been constructed for the vibrational analysis using OpenMolcas software [17].

**20.3 QM/MM Molecualr Dynamics**

Only B chain of tubulin was employed in this work. The model was solvated in a pre-equilibrated rectangular box of TIP3P [18] water with sodium ions added to neutralize the system. DFTB3 method [19] was used for the QM region. The QM region consists of the side chains of the Glu198 (protonated), Tyr200 and the entire PAC. Total number of atoms in QM region was calculated to be 74. For the MM region, Amber ff14SB force field [20] was used for all amino acids, and a second-generation of the GAFF force field (GAFF2) [21] was applied for the ligand. The cutoff radius value of QM area was 12 Å. For the system, one thousand minimization steps were performed, the temperature was slowly heated to 300K within 80 ps, followed by additional 1 ns equilibration and 4 ns DFTB3/MM MD simulation was carried out. The last frame was further subjected to a 60 ps B3LYP/MM MD with first 40 ps for equilibration and the rest 20 ps trajectory for checking the proton transfer between Oε atom of βGlu198 and N atom of PAC. The SHAKE algorithm [23] was applied to maintain all hydrogen atoms except for amino acid Glu198 during the B3LYP/MM MD production. In particular, B3LYP exchange correlation functional [22, 23] with a standard 6-31G basis set[24] was used for the QM region. The teraChem program [25], which was implemented in the Amber16 [26], was employed for DFT calculation at this stage.

**21. *In vivo* anti-cancer activity of PAC on mice models**

Animal Care and Use Committee of Sichuan University (Chengdu, Sichuan, China) approved the animal studies here. Five- to six-week-old female Balb/C athymic nude mice were obtained from HFK Bioscience Company（Beijing）. Mice continue to raise for 2 weeks to adapt to the environment, before implanted with 5🞨10^6^ Hct116 or H460 cells on the right flank. When the tumor volume reached about 100mm^3^, mice were divided into different groups and treated with 30mg/kg taxol (dissolved in physiological saline containing 10% ethanol and 10% tween-80) through intraperitoneal injection once a week or different dose of PAC (dissolved in physiological saline containing 5% ethanol and 5% tween-80) through tail vein injection every two days, tumor volume were recorded every two days and tumor volume were calculated as: 0.5 🞨length🞨width^2^.

**22. Statistical analysis**

Data are presented as mean±SD. Statistical differences were determined using an unpaired Student’s t test. p values are indicated in figure legend when necessary: **, p< 0.01; ***, p< 0.001.
